# Defining Early Steps in *B. subtilis* Biofilm Biosynthesis

**DOI:** 10.1101/2023.02.22.529487

**Authors:** Christine A. Arbour, Rupa Nagar, Hannah M. Bernstein, Soumi Ghosh, Yusra Al-Sammarraie, Helge C. Dorfmueller, Michael A. J. Ferguson, Nicola R. Stanley-Wall, Barbara Imperiali

## Abstract

The *Bacillus subtilis* extracellular biofilm matrix includes an exopolysaccharide that is critical for the architecture and function of the community. To date, our understanding of the biosynthetic machinery and the molecular composition of the exopolysaccharide of *B. subtilis* remains unclear and incomplete. This report presents synergistic biochemical and genetic studies built from a foundation of comparative sequence analyses targeted at elucidating the activities of the first two membrane-committed steps in the exopolysaccharide biosynthetic pathway. By taking this approach, we determined the nucleotide sugar donor and lipid-linked acceptor substrates for the first two enzymes in the *B. subtilis* biofilm exopolysaccharide biosynthetic pathway. EpsL catalyzes the first phosphoglycosyl transferase step using UDP-di-*N*-acetyl bacillosamine as phospho-sugar donor. EpsD is a GT-B fold glycosyl transferase that facilitates the second step in the pathway that utilizes the product of EpsL as an acceptor substrate and UDP-*N*-acetyl glucosamine as the sugar donor. Thus, the study defines the first two monosaccharides at the reducing end of the growing exopolysaccharide unit. In doing so we provide the first evidence of the presence of bacillosamine in an exopolysaccharide synthesized by a Gram-positive bacterium.

**IMPORTANCE:** Biofilms are the communal way of life that microbes adopt to increase survival. Key to our ability to systematically promote or ablate biofilm formation is a detailed understanding of the biofilm matrix macromolecules. Here we identify the first two essential steps in the *Bacillus subtilis* biofilm matrix exopolysaccharide synthesis pathway. Together our studies and approaches provide the foundation for the sequential characterization of the steps in exopolysaccharide biosynthesis, using prior steps to enable chemoenzymatic synthesis of the undecaprenol diphosphate-linked glycan substrates.

## INTRODUCTION

Biofilms are self-associating microbial systems that contain surface-adherent individuals within an extracellular matrix (1). The non-pathogenic bacterium, *Bacillus subtilis* (*Bs*), has been used extensively for understanding biofilm formation due to its ease of genetic manipulation and its extensive applied uses across diverse sectors of our economy (2). The *B. subtilis* biofilm matrix contains multiple specific components: BslA (a hydrophobin-like protein that confers hydrophobicity and structure to the community), fibers of the protein TasA (required for the structural integrity of biofilm), extracellular DNA (eDNA, important at early stages of biofilm formation), poly-γ-glutamic acid (γ-PGA, possible function in water retention), and an exopolysaccharide (EPS) (3).

The EPS is the main carbohydrate component of the *B. subtilis* matrix and is critical for biofilm architecture and biofilm function (4, 5). Despite considerable interest in understanding biofilm biosynthesis and regulation, the individual building blocks for this macromolecular glycoconjugate have not been determined. Biosynthesis of EPS is dependent on enzymes expressed from a 15-gene *epsABCDEFGHIJKLMNO* (*epsA–O*) operon, which has a similarity with the *Campylobacter jejuni pgl* operon (**Figure 1A**) (6). These enzymes have been annotated based on sequence analysis as a phosphoglycosyl transferase (PGT), glycosyl transferases (GTs), uridine diphosphate sugar (UDP-sugar) modifiers, a regulatory enzyme, and a flippase (5, 7, 8). However, most of the membrane-associated enzymes that are involved in the biosynthesis of exopolysaccharide in *B. subtilis* have not been biochemically characterized. Furthermore, analysis of exopolysaccharide composition has afforded conflicting information. Even studies of the same strain of *B. subtilis* (namely NCIB 3610) provided different carbohydrate compositions depending on the bacterial growth conditions and/or methods of extraction and purification (6). For example, when grown in a glutamic acid and glycerol-rich media, an EPS fraction contained glucose, *N*-acetylgalactosamine (GalNAc), and galactose (Gal) (9, 10). The same strain grown in lysogeny broth (LB) media that included magnesium and manganese divalent cations, produced an EPS fraction containing mannose and glucose (11, 12). Furthermore, growth in a minimal media supplemented with glucose (MMG) produced an EPS fraction containing poly-*N*-acetylglucosamine (GlcNAc) (5).

**Figure 1.**
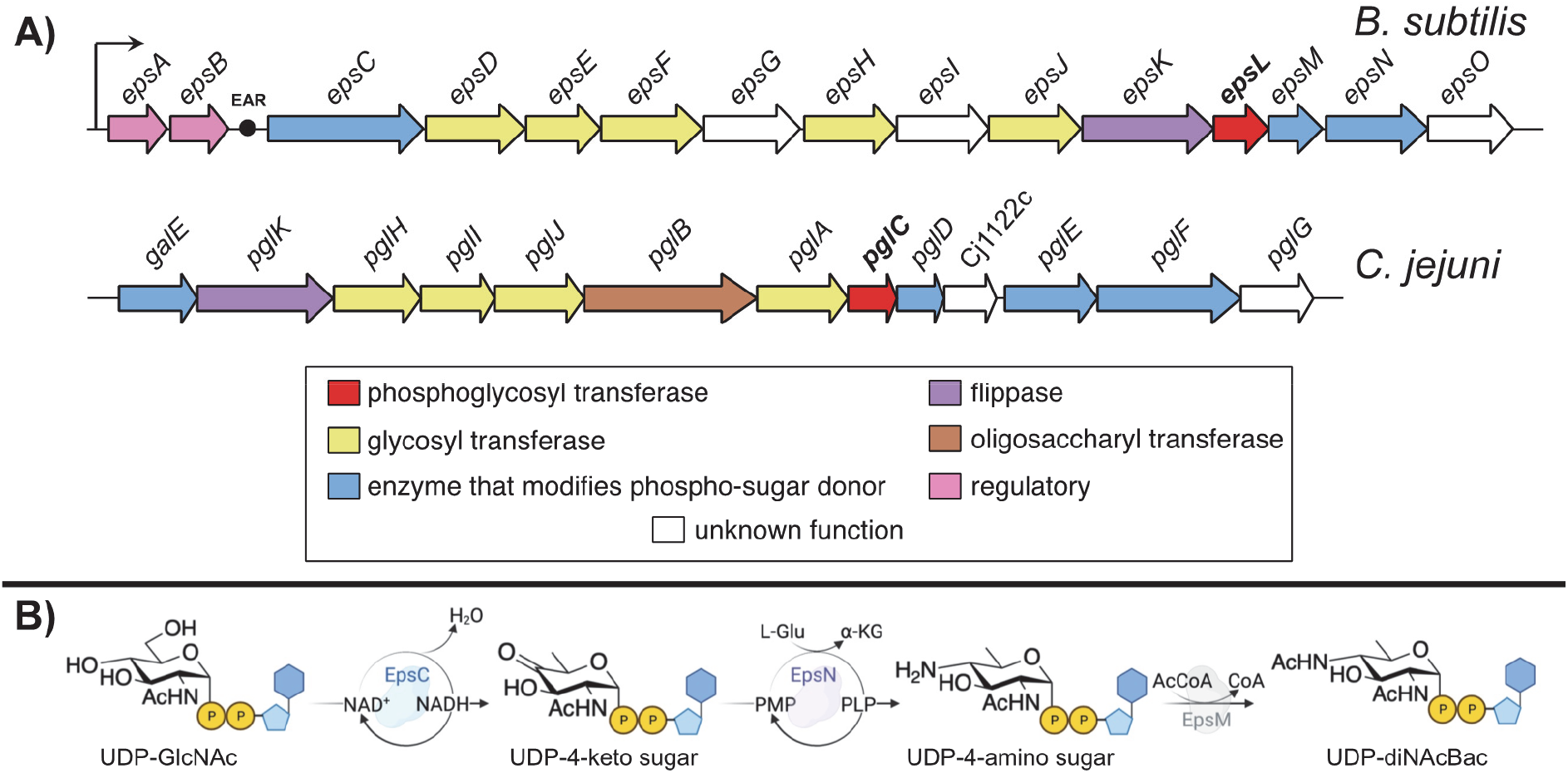
Comparison of glycoconjugate synthesis in *B. subtilis* and *C. jejuni*. **A)** The *epsA–O* operon of *B. subtilis* and the *pgl* operon of *C. jejuni* drawn broadly to scale. EAR represents the *eps*-associated RNA (21) situated between *epsB* and *epsC*. **B)** The biosynthesis of UDP-diNAcBac in *B. subtilis* catalyzed by EpsCNM.

UDP-*N,N*’-diacetylbacillosamine (UDP-diNAcBac) is a prokaryote-specific nucleotide sugar donor (13). The monosaccharide component, diNAcBac, was originally discovered in *Bacillus licheniformis* (14). Based on *in vitro* activity and sequence similarity, EpsC, EpsN, and EpsM are proposed to produce UDP-diNAcBac in *B. subtilis* (**Figure 1B**). EpsC contains the sequence motifs found in other dehydratases and is a UDP-GlcNAc 4,6-dehydratase that converts UDP-*N*-acetylglucosamine (UDP-GlcNAc) to UDP-2,6-dideoxy-2-acetamido 4-keto glucose (UDP-4-keto) (15). It catalyzes the NAD^+^-dependent elimination of water across C5 and C6, while oxidizing C4 of UDP-GlcNAc. The resulting α,β-unsaturated ketone is reduced by hydride addition at C6, followed by tautomerization and regeneration of NAD^+^ to provide the UDP-4-keto sugar and cofactor for a new catalytic cycle. The penultimate enzyme, EpsN, is a pyridoxal 5’-phosphate (PLP)-dependent aminotransferase that transfers an amine from L-glutamate to the C4 of UDP-4-ketosugar to provide UDP-2,6-dideoxy 2-acetamido 4-amino glucose (UDP-4-amino) (16). The subsequent enzyme, EpsM, is an acetyltransferase that transfers an acetyl group from acetyl coenzyme A (AcCoA) onto UDP-4-amino sugar to provide UDP-diNAcBac (17). To further support the assignment of these Eps enzymes, isofunctional homologs in *Campylobacter*, in particular *C. jejuni* (PglF, PglE, PglD) (**Figure 1A**), have been biochemically characterized and shown to make UDP-diNAcBac in a similar fashion (13, 18-20). Consistent with this, EpsCNM from *B. subtilis* and PglFED from *C. jejuni* (*Cj*) have 54, 64, and 50% sequence similarity, respectively (15).

Our overarching goal is to elucidate the composition and structure of the *B. subtilis* biofilm matrix EPS. Given the inconsistencies obtained from direct analysis of the extracted EPS material, we elected to start by determining the identity of the individual monosaccharides at the reducing end of the exopolysaccharide. In this work, we investigate and define the substrate specificity of two enzymes encoded within the *eps* operon, EpsL and EpsD, annotated as a phosphoglycosyl transferase and glycosyl transferase, respectively, using biochemical and genetic complementation approaches. We present experimental evidence supporting the designation of EpsL as a PGT which installs diNAcBac as the first monosaccharide onto a undecaprenol phosphate (UndP) carrier. We also identify EpsD as the second enzyme, and the first GT, in the pathway that likely installs GlcNAc onto the diNAcBac-appended lipid anchor. Thus, a key polyprenol-diphosphate-linked disaccharide is proposed and can be made available through chemoenzymatic synthesis. Therefore, our work sets the stage for future analysis of downstream glycosyltransferase reactions in the EPS pathway.

## Results

### Characterizing the phosphoglycosyl transferase (EpsL) in the EPS biosynthetic pathway

Phosphoglycosyl transferases (PGTs) are enzymes responsible for catalyzing the first membrane-committed step in many essential glycosylation pathways by transferring a sugar-phosphate onto a lipid acceptor carrier. PGTs are represented by two distinct membrane topologies, mono-and polytopic, (22) and perform mechanistically-distinct modes of catalysis (23). The monoPGTs comprise three families: small, long, and bifunctional enzymes. The sequence similarity network (SSN) of small monoPGTs provided an uncharacterized enzyme from *B. subtilis*, EpsL (24). *B. subtilis* EpsL contains the key residues that are the hallmarks of the monoPGTs catalytic domain and other signature motifs (**Figure 2**) (25). These include a basic (KR) motif near the N-terminus and helix-break-helix (SP) motif in the membrane-associated domain that contribute to the membrane reentrant topology of the enzyme. Additionally, the catalytic dyad (DE) that is responsible for covalent catalysis, and the uridine-binding residues (PRP) are present. Furthermore, EpsL is similar to small monoPGTs from other Gram-positive bacteria (*Staphylococcus aureus* (*Sa*) 42% identity), a PGT that has been shown to use UDP-D-FucNAc as the sugar-phosphate donor substrate (26). However, higher sequence similarity is observed with PglCs from *campylobacter* (*C. concisus (Cc)* 58%,*C. jejuni (Cj)* 59% identity) and *Helicobacter pullorum* (*Hp*) (60% identity) (**Figure 2**). Based on sequence similarity with monoPGTs from *C. concisus* and *C. jejuni*, we hypothesized that EpsL uses UDP-diNAcBac. This is consistent with the conclusion that EpsCNM synthesize this particular UDP-sugar (15-17).

**Figure 2.**
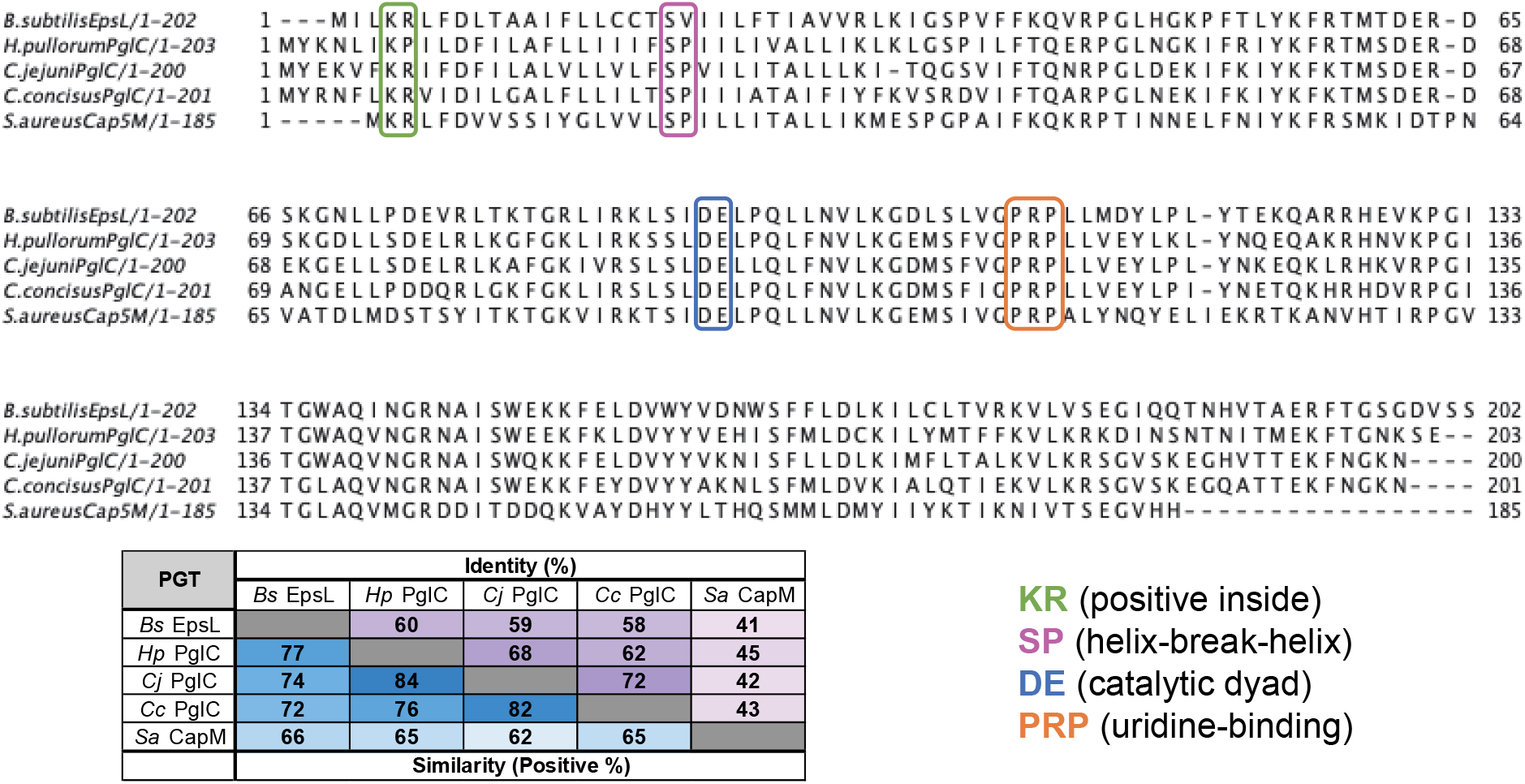
Protein sequence comparison of select monotopic phosphoglycosyl transferases (monoPGTs). Sequence alignment of *Bs* EpsL with monoPGTs from Gram-positive and Gram-negative bacteria made in Jalview.(27, 28) The basic local alignment search tool (BLAST) was used to obtain percent identity and similarity from accession numbers: *Bs* EpsL (P71062), *Hp* PglC (E1B268), *Cj* PglC (Q0P9D0), *Cc* PglC (A7ZET4), *Sa* CapM (P95706) with more details in the supporting information (**Table S1**).

### Biochemical and genetic evaluation of EpsL substrate specificity

To test the hypothesis that EpsL uses UDP-diNAcBac as the phospho-sugar donor substrate, heterologous expression of *epsL* was carried out in *E. coli* following a previously described protocol for monotopic PGTs from *C. concisus* and *C. jejuni* (23, 29, 30). After isolation of the cell envelope fraction, eight detergents were screened to evaluate the solubilization efficiency and purity of the enzyme (**Figure S1**) (**Table S2**). The detergent solubilization screen provided two detergents, Triton X-100 and octaethylene glycol monododecyl ether (C_12_E_8_), that efficiently solubilized EpsL, while minimizing the solubilization of undesired proteins from the cell envelope fraction. For that reason, EpsL was solubilized and purified in Triton X-100 and C_12_E_8_ on a preparative scale for downstream applications (**Figure 3A**) (**Figure S2**).

**Figure 3.**
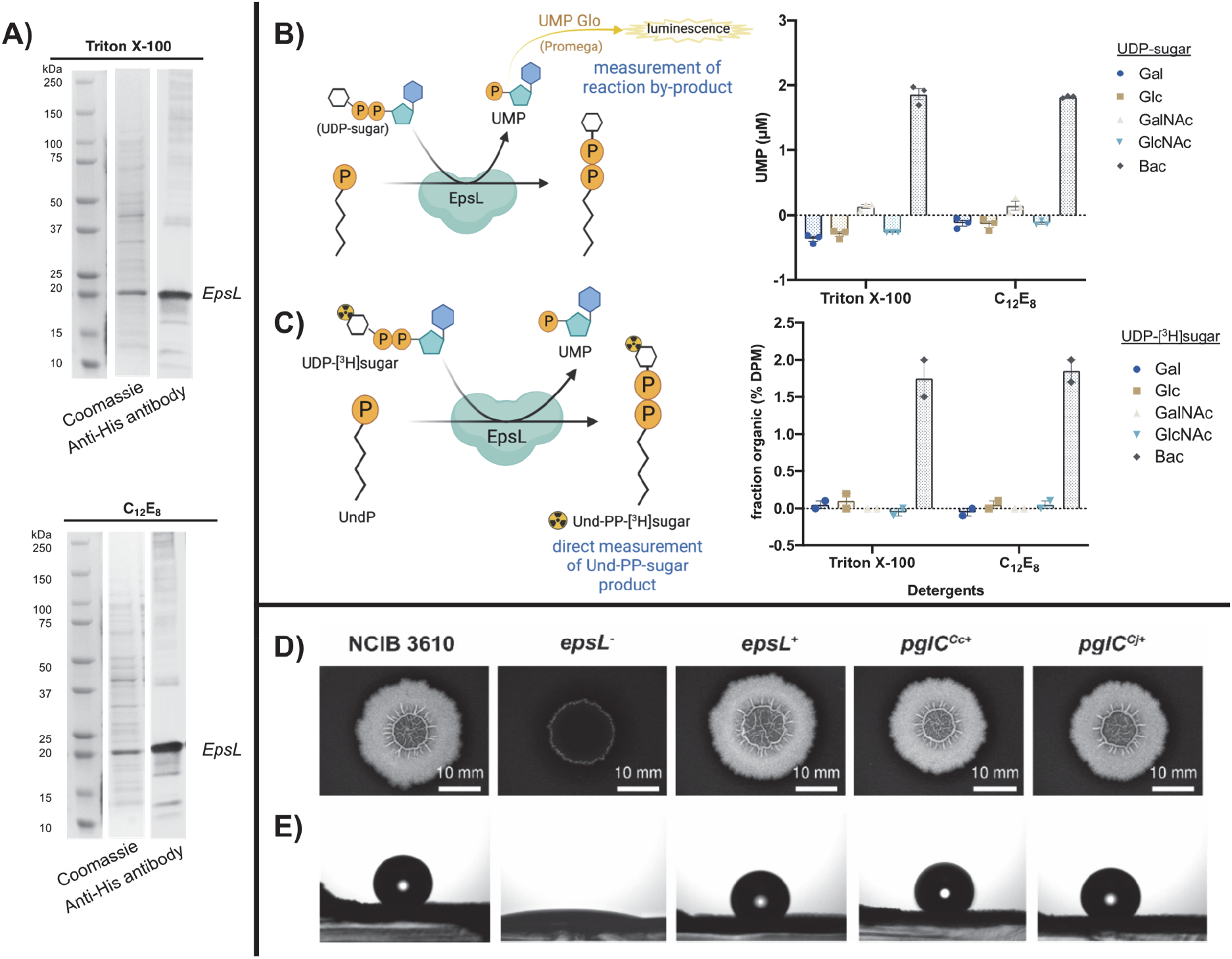
Purification and biochemical and phenotypic characterization of EpsL. **A)** *B. subtilis* EpsL purification visualized by SDS-PAGE (Coomassie) and anti-His antibody western blots. **B**) Complementary biochemical activity assays of *B. subtilis* EpsL. A luminescence-based assay, UMP Glo, which measured the UMP byproduct of the PGT reaction. Error bars are given for mean ± SEM, n = 3. **C)** A radioactivity-based assay that measures the Und-PP-[^3^H]sugar product. Error bars are given for mean ± SEM, n = 2. **D)** and **E**) Genetic complementation of *ΔepsL-Bs* mutant with *pglC* of *Campylobacter*. **D)** represents colony biofilm morphologies of wild-type (*B. subtilis* NCIB 3610) *ΔepsL* mutant (*epsL*^-^ - NRS5907) and genetically complemented strains (*epsL*^+^ - NRS5942, *pglC*^*Cc*+^ - NRS6692, *pglC*^*Cj*+^ - NRS6618, see **Table S3**). The colony biofilms were grown at 30 °C for 48 hours prior to imaging. **E)** represents the respective sessile water drop analysis of the colony biofilms with a 5 μl water droplet on top. The representative images were taken after 5 min, except *epsL*^-^ where the image was taken at 0 min due to extreme hydrophilicity of the surface in absence of biofilm.

The activity of solubilized and purified EpsL was evaluated. This was achieved through a substrate screen with five UDP-sugar donors and UndP as a lipid acceptor using two complementary biochemical assays; UMP Glo® and a radioactivity-based assay (**Figure 3B**-**C**). The standard commercial ^3^H-labeled and unlabeled UDP-sugars (UDP-Gal, UDP-Glc, UDP-GalNAc, and UDP-GlcNAc) were used for the screens (**Figure S3**). Additionally, UDP-diNAcBac and UDP-[^3^H]diNAcBac, both prepared via chemoenzymatic methods, were used (**Scheme S1**) (**Figure S4**). The UMP Glo® assay developed by Promega monitors the production of UMP over the course of a reaction (**Figure 3B**) (31). This indirect measurement of reaction progress is excellent for initial screens of PGTs. However, to quantify the reaction more directly, an assay that monitors the reaction product is needed. For that reason, we employed a radioactivity-based assay to directly measure the formation of the Und-PP-sugar following liquid-liquid extraction of the Und-PP-linked product (**Figure 3C**) (**Figure S5**). We additionally monitored reaction progress in non-radioactive reactions by normal phase silica thin layer chromatography (TLC) **Figure S6**). During the reaction, a new product was formed that had the same retention factor (R_f_) as the authentic standard Und-PP-diNAcBac from *C. concisus* PglC (32) providing biochemical evidence that EpsL can use UDP-diNAcBac as donor substrate in the presence of the UndP acceptor.

We proposed that if EpsL was a PGT that installs diNAcBac as the first monosaccharide in the EPS pathway, then PglC of *Campylobacter* should be able to substitute for EpsL activity *in vivo*. In the absence of *epsL, B. subtilis* is unable to form the rugose, hydrophobic colony biofilms on agar plates typical of those formed by strain NCIB 3610 (**Figure 3D**). Therefore the *B. subtilis epsL* deletion strain was genetically complemented with the PGT coding sequences from *C. jejuni* and *C. concisus* (PglC) (**Table S3-6**). The coding sequences were placed under the control of an IPTG inducible promoter and integrated into the chromosome at the ectopic *amyE* gene in the *epsL* deletion strain. The *B. subtilis epsL* coding region was used as a positive control (see **Table S3**) (**Figure S7**). In each case, in the presence of 25 μM IPTG, the genetic complementation of the *epsL* deletion strain by the *pglC* coding region was noted. The presence of *pglC* provided full recovery of the rugose colony biofilm architecture to the *epsL* deletion strain (**Figure 3D**). Additionally, recovery of both the area occupied by the mature colony biofilm (**Figure S7B**) and surface hydrophobicity (**Figure 3E**) (**Figure S7C**) was observed to a level that was indistinguishable from the analysis of the NCIB 3610 parental strain. Taken together with the bioinformatic analysis, our biochemical and genetic data support the designation of EpsL as a PGT that installs diNAcBac as the first monosaccharide at the reducing end of the *B. subtilis* EPS.

### Substrate specificity of EpsD, the first glycosyl transferase in the EPS pathway

By determining the first membrane-committed step in the EPS pathway, we were provided with an experimental system where we could use the product of EpsL (Und-PP-diNAcBac) to study the first glycosyl transferase in the pathway. As the structures of glycosyl transferases are relatively similar it is not possible to predict the substrate specificity from sequence alone. In the *Campylobacter pgl* pathways, the PglA enzyme is responsible for the second step in the glycan biosynthetic pathway, catalyzing the transfer of GalNAc from UDP-GalNAc to Und-PP-diNAcBac (32). There are five GTs encoded by the *epsA-O* operon: EpsD, EpsE, EpsF, EpsH, and EpsJ (**Figure 1A)**. Of these, EpsD and EpsFare the most similar to PglA at the sequence level (**Figure 4A**) (**Table S7**). Both belong to GT-4 family in the CAZy classification (33) and AlphaFold structural analysis suggests that both possess a GT-B fold, like PglA. In contrast, the remaining GTs encoded by *epsA-O* operon, EpsE, EpsH and EpsJ, belong to GT-2 family and are predicted to have GT-A folds. Therefore, based on the sequence similarities of EpsD and EpsF to PglA and their GT structural fold analyses, we predicted either EpsD or EpsF could be the first glycosyltransferase in EPS pathway.

**Figure 4.**
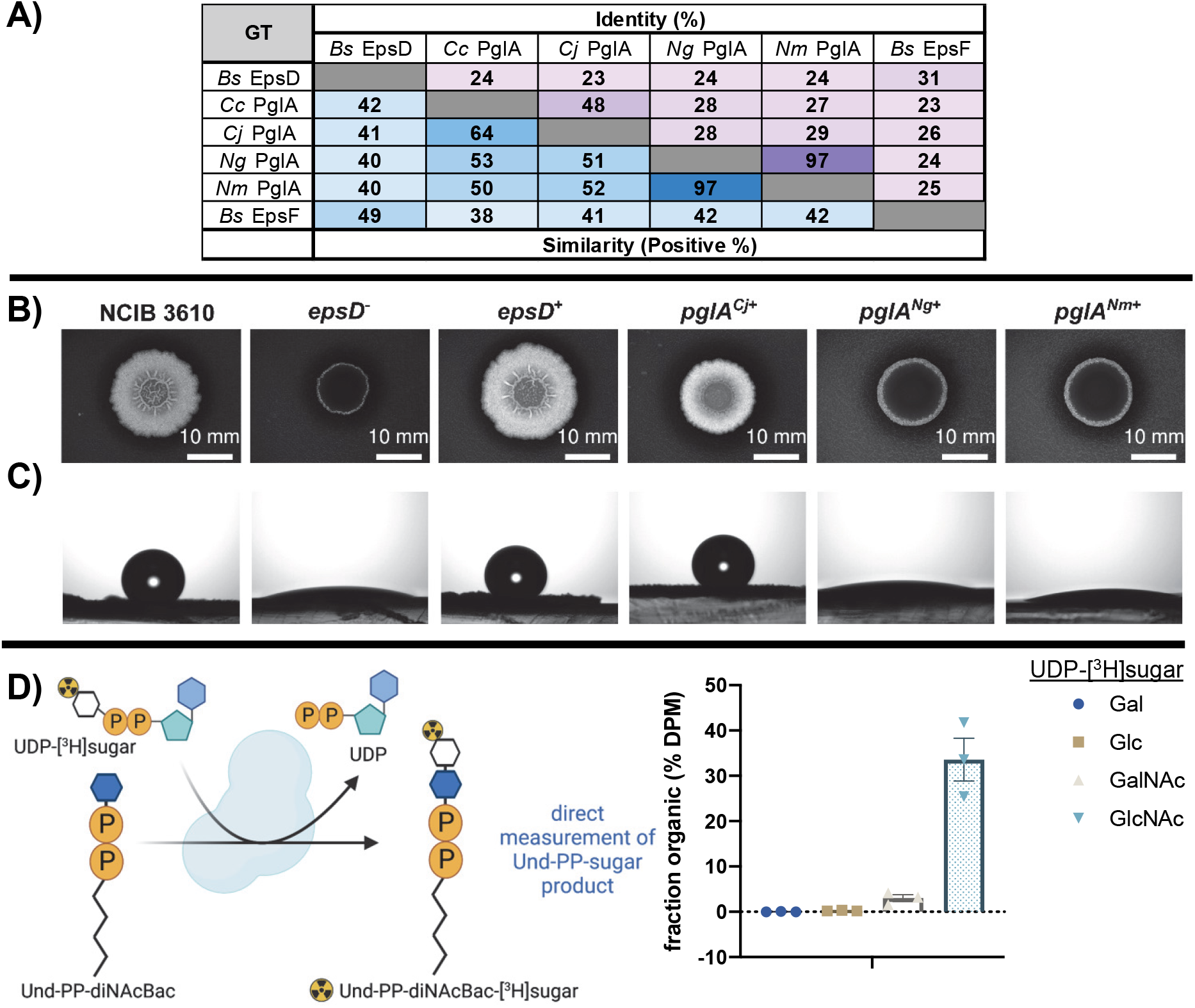
Sequence comparison and biochemical and phenotypic analysis of EpsD. **A)** Sequence identity of *B. subtilis* EpsD with characterized PglAs from Gram-negative bacteria. Accession numbers: *Bs* EpsD (P71053), *Cc* PglA (A7ZET5), *Cj* PglA (A0A2U0QT38), *Ng* PglA (Q5F602), *Nm* PglA (Q9K1D9), and *Bs* EpsF (P71055). **B) and C)** Genetic complementation of *Bs ΔepsD* mutant with *pglA* of *Campylobacter* and Neisseria. **B)** represents colony biofilm morphologies of wild-type (*B. subtilis* NCIB 3610), *ΔepsD* mutant (*epsD*^-^ - NRS5905) and genetically complemented strains (*epsD*^+^ - NRS5930, *pglA*^*Cj*+^ - NRS6605, *pglA*^*Ng*+^ - NRS6619, *pglA*^*Nm*+^ - NRS6620). The colony biofilms were grown at 30°C for 48 hours prior to imaging. **C)** represents the respective sessile water drop analysis of the colony biofilms with a 5 μl water droplet on top. The representative images of wild-type, *epsD*^+^ and *pglA*^*Cj*+^ were taken after 5 min, whereas the images of *epsD*^-^ mutant, *pglA*^*Ng*+^ and *pglA*^*Nm*+^ were taken at 0 min due to extreme hydrophilicity of the surface in absence of biofilm. **D)** Biochemical determination of substrate specificity of *Bs* EpsD with Und-PP-diNAcBac as an acceptor substrate in a radioactive-based assy. Error bars are given for mean ± SEM, n = 3.

Based on the hypothesis that EpsD or EpsF could be the PglA homologue in *B. subtilis* (**Figure 4A**), we tested whether PglA could functionally substitute for either EpsD or EpsF *in vivo*. We therefore investigated the genetic complementation *B. subtilis epsD* and *epsF* deletion strains by the PglA coding sequences from *C. jejuni* and other related UDP-Gal transferase enzymes from *Neisseria gonorrhoeae* (*Ng*) and *Neisseria meningitidis* (*Nm*) (**Figure 4B**) (**Figure S8A**). The *epsD* and *epsF* deletion strains of *B. subtilis* are unable to form the wild-type rugose, hydrophobic colony biofilms on agar plates (**Figure 4B**) (**Figure S8A**). The *pglA* genes were placed under the control of an IPTG inducible promoter and integrated into the chromosome at the ectopic *amyE* gene in the *epsD* and *epsF* deletion strains. The *B. subtilis epsD* and *epsF* coding regions were used as the respective positive controls (see **Table S3**), (**Figure 4B**), (**Figure S8A-B**). In the presence of 25 μM IPTG, the genetic complementation of the *epsD* deletion strain by *pglA* gene of *C. jejuni* resulted in partial recovery of biofilm formation, whereas complementation with *pglA* genes from *Neisseria* did not recover the biofilm phenotype (**Figure 4B**). In the case of *pglA from C. jejuni*, in addition to a partial rescue of biofilm architecture, there was recovery of both the area occupied by the mature colony biofilm (**Figure S8C**) and surface hydrophobicity (**Figure 4C**) (**Figure S8D**). The measurements quantified in each case were indistinguishable from those obtained from the analysis of the NCIB 3610 parental strain. In contrast, although the *epsF* deletion strain could be fully complemented by the *epsF* coding region, expression of the *pglA* genes were unable to recover the biofilm formation (**Figure S8A**). This conclusion is supported by AlphaFold modeling of the *Bs* EpsD, EpsF and *Cj* PglA structures where EpsD and PglA (rsmd 1.4Å) share an overall higher structural similarity than EpsF and PglA (rsmd 4.1Å) (**Figure S9**).

We next took a biochemical approach to confirm the activity of EpsD by using the purified Und-PP-diNAcBac from chemoenzymatic synthesis. To investigate the identity of the UDP-sugar donor for EpsD we used heterologous expression of EpsD in *E. coli* (**Figure S10**). Initial attempts to detergent solubilize EpsD were made and provided protein as assessed by SDS-PAGE (**Figure S11**). However, the enzyme was no longer active once solubilized from the cell envelope fraction (Arbour, Bernstein, Ghosh, Imperiali, unpublished data). For that reason, we determined the UDP-sugar substrate using the cell envelope fraction of *E. coli* expressing the *epsD* using a radioactivity-based assay with Und-PP-diNAcBac, the substrate produced from EpsL (**Figure 4D**) (**Figure S12**). The panel of donor substrates used for the activity assay were tritiated, commercially-available UDP-[^3^H]sugars, namely UDP-[^3^H]Gal, UDP-[^3^H]Glc, UDP-[^3^H]GalNAc, and UDP-[^3^H]GlcNAc. We determined that in the presence of UDP-[^3^H]GlcNAc, EpsD converts 35% of the total amount of UDP-[^3^H]GlcNAc to Und-PP-diNAcBac-[^3^H]GlcNAc. Additionally, under identical conditions, low transfer (4%) of [^3^H]GalNAc to Und-PP-diNAcBac-[^3^H]GalNAc was observed (**Figure 4D**). Therefore, we conclude that EpsD can use Und-PP-diNAcBac as an acceptor substrate for the transfer of GlcNAc. Regarding the stereochemistry of the new glycosidic linkage, we examined the sequences of EpsD with PglA from *C. jejuni* and *C. concisus* and the structural overlay of AlphaFold models of EpsD and PglA (*C. concisus*) (**Figure S13**). These analyses strongly suggest that EpsD follows a similar mechanistic course affording an α-1,3-linkage, which is achieved through a retaining GT mechanism.(34) Additionally, EpsD displays substrate promiscuity by accepting UDP-GalNAc as a less preferred substrate (**Figure 4**).

## Discussion

It is extremely challenging to elucidate the structures of complex glycoconjugates directly from bacterial extracts. A case in point is the major polysaccharide found in the extracellular matrix of *B. subtilis* biofilms, which has remained undefined, despite considerable experimentation for many years. This is an important area of research as biofilm formation is a prevalent behavior displayed across multiple microbial species and exopolysaccharide production is highly correlated with biofilm formation (35). In this study, we have applied complementary biochemical and genetic approaches to establish the function of essential enzymes that catalyze key early steps in biofilm biosynthesis from the *Bacillus subtilis epsA-O* operon. Overall, the sequences of protein encoded by the operon support the expression of enzymes involved in UDP-sugar biosynthesis as well as several GTs and a PGT with unknown substrate specificity and roles in biofilm biosynthesis, however, in the absence of targeted analysis, the eps pathway cannot be defined.

### EpsL is a functional PGT that utilizes UDP-diNAcBac

Bioinformatic analysis suggested that many of the genes in the *epsA-O* cluster showed similarity to the *pgl* gene cluster, which is responsible for the general protein N-glycosylation pathway in *C. jejuni* (32, 36). As the *pgl* gene cluster had been biochemically characterized and shown to be involved in the biosynthesis of UDP-diNAcBac and a heptasaccharide product containing diNAcBac at the reducing end of the glycan (37), this similarity provided the foundation for exploration of the function of selected enzymes in the *B. subtilis* EPS pathway. Previous sequence analysis and *in vitro* characterization of EpsCNM suggested that these enzymes are responsible for the biosynthesis of UDP-diNAcBac (15-17). Sequence analysis also identified EpsL as a close homolog of the *C. jejuni* and *C. concisus* PglCs, which are structurally and biochemically well-characterized PGTs (**Figure 2**) (32). The identification of a PGT is noteworthy as these enzymes catalyze phosphosugar transfer from UDP-diNAcBac to a polyprenol phosphate carrier as the first membrane-associated step in many glycoconjugate assembly pathways (38).

Thus, we designed a strategy to implement an *in vitro* biochemical activity assay using UndP as the acceptor substrate and a series of [^3^H]-labeled and unlabeled UDP-sugars, including UDP-diNAcBac. Following heterologous expression, solubilization, and purification, EpsL was used to screen enzyme activity *in vitro*. Complementary assays using either radiolabeled sugars or the UMP-Glo® assay were applied to confirm that EpsL prefers UDP-diNAcBac as phosphosugar donor and affords the Und-PP-diNAcBac product (**Figure 3B**-**C**). These *in vitro* biochemical assay results were supported by genetic analyses using biofilm formation as the phenotypic readout. This revealed that the *B. subtilis epsL* deletion mutant could be genetically complemented by the *pglC* coding sequence of *C. jejuni* (**Figure 3D**). Thus, we conclude that EpsL catalyzes the first step in EPS biosynthesis pathway to form Und-PP-diNAcBac. Moreover, we show the first experimental evidence of the function of a UDP-diNAcBac utilizing PGT in a Gram-positive bacterium and the presence of diNAcBac as the first sugar at the reducing end of EPS in *B. subtilis*. These findings are significant; diNAcBac was first discovered in *Bacillus licheniformis* (14), however, to date the diNAcBac sugar has only been described in N- and O-linked glycoproteins, lipopolysaccharide (LPS), and the capsular polysaccharide (CPS) of diverse Gram-negative bacteria (13).

### EpsD is a UDP-GlcNAc-dependent *N*-acetyl glucosamine transferase in *B. subtilis*

The successful characterization of the first step in EPS pathway provided the Und-PP-diNAcBac substrate for exploring the next enzyme in the EPS biosynthesis. In this case, although the *epsA-O* gene cluster revealed five candidate GTs with predicted GT-A or GT-B fold, the assignment of structure to functional specificity could not be definitively predicted. However, the similarity of *epsA-O* cluster genes with *C. jejuni* N-glycosylation pathway genes helped us to narrow down the candidates to EpsD and EpsF as possible GTs for the subsequent step in the pathway. Our bioinformatic analysis suggested that both EpsD and EpsF share similarity with PglA of *C. jejuni* and selected *Neisseria* sp. (**Figure 4A**) and we additionally knew that both EpsF and EpsD were essential for biofilm formation in *B. subtilis* (5). The possibility that EpsF was the next enzyme in the biosynthetic pathway was ruled out by the inability of *pglA* genes of *C. jejuni* and *Neisseria* sp. to rescue the biofilm formation upon expression in *epsF* deletion mutant of *B. subtilis* (**Figure S8A**). However, comparable experiments with EpsD provided new insight as genetic complementation with the *C. jejuni pglA* was able to partially rescue the biofilm-negative phenotype in the *epsD* deletion mutant of *B. subtilis* (**Figure 4B**). In contrast, the expression of two *pglA variants*, which catalyze the addition of Gal in the second step of the *Neisseria* pgl pathway(39) did not rescue the phenotype in the *epsD* deletion mutant. Although, the partial complementation of p*glA* of *C. jejuni* in *epsD* deletion mutant did not confirm the preference of EpsD for GalNAc it provided the possibility that the preferred sugar substrate could be the related HexNAc sugar, GlcNAc. This hypothesis was supported by using a biochemical approach where the cell envelope fraction of *E. coli* expressing EpsD was used to assess activity using Und-PP-diNAcBac and four different commercially available ^3^H-labeled UDP-sugars as donor substrates. The *in vitro* assay results provided further insight into the EpsD sugar substrate selectivity; EpsD showed a clear preference for UDP-GlcNAc over the other UDP-sugars tested with significant conversion UDP-[^3^H]GlcNAc to Und-PP-diNAcBac-[^3^H]GlcNAc (**Figure 4D**). This supports the function of EpsD in the second step in the EPS pathway. Interestingly, EpsD was also able to transfer [^3^H]GalNAc to Und-PP-diNAcBac although with far lower efficiency. This donor substrate promiscuity displayed by EpsD not only explains the partial genetic complementation of *epsD* deletion mutant of *B. subtilis* with p*glA* of *C. jejuni* but also provides insight into the step downstream. As previously established, PglA transfers GalNAc onto Und-PP-diNAcBac in *C. jejuni* N-glycans (32, 40). Thus the partial complementation observed upon expressing *pglA* in the *B. subtilis epsD* deletion mutant suggests that Und-PP-diNAcBac-GalNAc is not a preferred acceptor for the next GT in the*B. subtilis* EPS biosynthetic pathway, resulting in the observed partial biofilm phenotype. It also suggests possible acceptor substrate promiscuity of the next GT in line.

### Summarizing new insights into the *Bacillus subtilis* EPS biosynthetic pathway

The characterization of EpsL and EpsD in this study has set the foundation for characterizing the remaining GTs in the EPS biosynthesis pathway, which would ultimately enable us to define the EPS sugar composition and structure. Based on the experimental evidence provided in this study, we propose the current EPS glycosylation pathway (**Figure 5**). EpsCNM have already been shown to biosynthesize UDP-diNAcBac (15-17). EpsL is a PGT that transfers diNAcBac onto Und-P converting it to Und-PP-diNAcBac. EpsD further extends this glycan by transferring GlcNAc onto the product from EpsL thus converting it to Und-PP-diNAcBac-GlcNAc. These findings also indicate a divergence in the *B. subtilis* EPS glycosylation pathway after the synthesis of Und-PP-diNAcBac (as diNAcBac-GlcNAc-) compared to *C. jejuni* (diNAcBac-GalNAc-) and *N. gonorrhoeae* (diNAcBac-Gal-) pathways. Homologs of EpsL and EpsD are present broadly across the *B. subtilis* clade. This suggests the presence of similar glycosylation pathways and exopolysaccharides in many *Bacillus* species and provides an opportunity to explore the diversity of diNAcBac-containing clusters and the associated exopolysaccharides.

**Figure 5.**
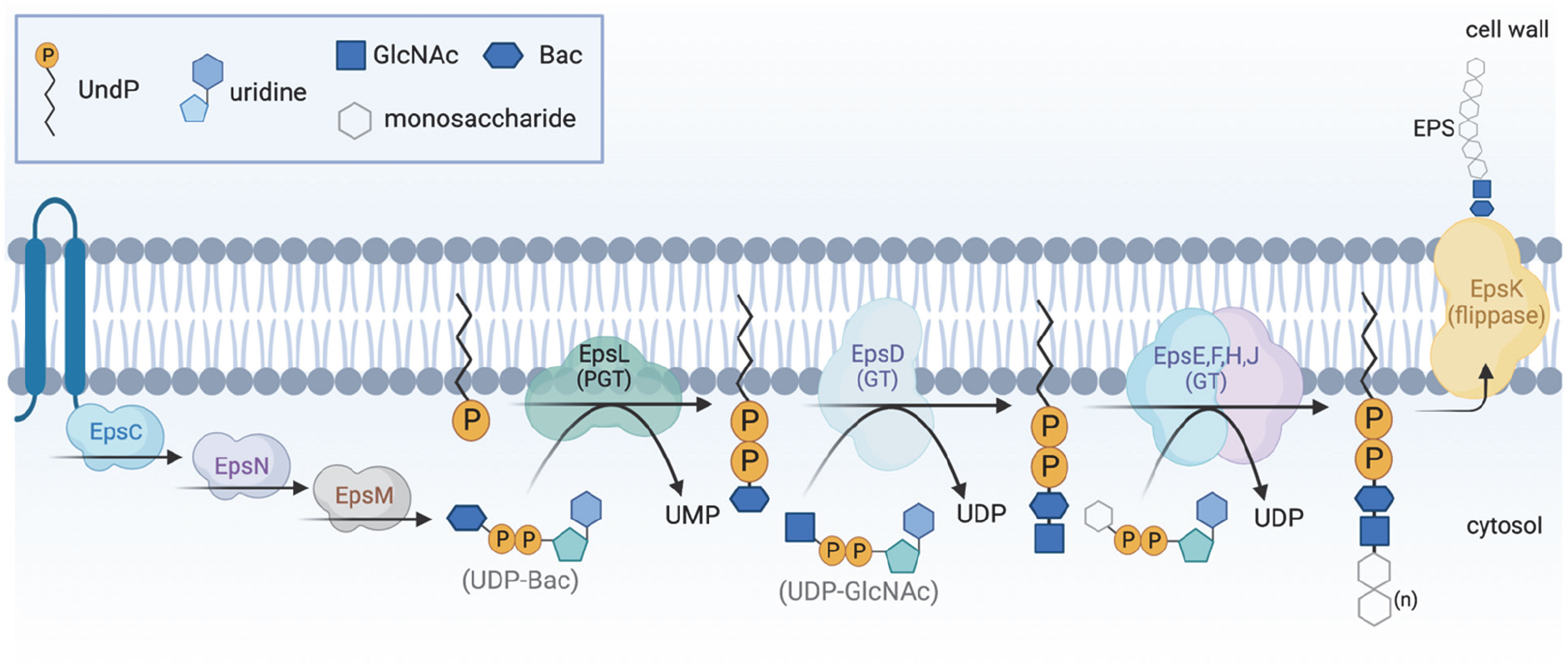
The proposed biofilm matrix exopolysaccharide biosynthetic pathway in *B. subtilis*. EpsCNM synthesize UDP-diNAcBac, which severs as a donor substrate for EpsL. EpsL transfers diNAcBac onto Und-P and EpsD catalyzes the second step and transfers GlcNAc from a UDP-GlcNAc sugar donor. The next GTs functioning downstream are to be characterized.

### Overarching Conclusion

The study of glycoconjugate biosynthesis pathways requires a concerted effort of different approaches as individual bioinformatic, biochemical, and genetic approaches often provide incomplete details. In this study, we establish the sequential characterization of the *B. subtilis* EPS steps by applying biochemical assays and phenotypic screening to the first two membrane-associated processes in the pathway – EpsL and EpsD. The major advantage of addressing steps in the pathway in their biosynthetic order is that the characterization of each enzyme provides the substrate for investigating the following step. Additionally, as enzyme expression and isolation (either in a cell envelope fraction or in a detergent-solubilized form) is included in the process, it enables the chemoenzymatic synthesis of products for additional analysis and use in related pathways. The established enzyme assays also provide the opportunity for small molecule inhibitor screening, both individually (EpsL or EpsD) or as biosynthetic partners (EpsL and EpsD). Taken together, these studies set a clear course for analysis of the downstream EPS glycosylation pathway and the development of a complete picture of EPS structure.

## Acknowledgements

We thank Natalie Bamford for reading the manuscript and providing helpful input. The work at the University of Dundee was funded by the Biotechnology and Biological Science Research Council (BBSRC) [BB/P001335/1, BB/R012415/1]. We thank the National Institute of Health (NIH) for financial support to B.I. (GM039334 and GM131627) and C.A.A. (F32GM136023).

**Figures 1B, Figure 3B-C, Figure 4D** and **Figure 5** were created with BioRender.com.

## Credit Statement

C.A.A., H.M.B., R.N., N.S.W., H.D. M.A.J.F., Y.A.S., B.I. designed this study, C.A.A., R.N., S.G., Y.A.S., H.M.B. collected the data, and all authors interpreted the data. All authors were involved in the writing and editing of the manuscript.

