## Supplemental Information for "Defining Early Steps in *B. subtilis* Biofilm Biosynthesis"

### Table of Contents

General Information ........ S3

**Table S1.** UniProt accession numbers and associated protein sequences of monoPGTs used for sequence identity ........ S3

Overexpression of *Bs* EpsL and *Bs* EpsD ........ S4

Detergent screening of *Bs* EpsL ........ S5

**Table S2.** The concentration and structures of the detergents used in the solubilization screen ........ S5

**Figure S1.** *Bs* EpsL detergent solubilization screen gels ........ S6

**Figure S2.** *Bs* EpsL detergent solubilization with Triton X-100 and $C_{12}E_8$ gels ........ S7

**Figure S3.** Uridine diphosphate sugar structures used in this study ........ S8

Protein expression for UDP-diNAcBac chemoenzymatic synthesis ........ S8

Western blotting analysis ........ S8

UDP-diNAcBac chemoenzymatic synthesis ........ S10

**Scheme S1.** Chemoenzymatic synthesis of UDP-diNAcBac ........ S10

**Figure S4.** TLC of the chemical acetylation reaction of UDP-4-amino sugar .................................. S11

UDP-[$^3$H]diNAcBac enzymatic synthesis and purification .......................................... S12

Radioactivity-based biochemical assays on purified protein (*Bs* EpsL) ...................... S12

**Figure S5.** Substrate specificity determination with *B. subtilis* EpsL using a radioactivity-based assay ........................................................................................................ S13

UMP Glo biochemical assays ........................................................................................ S13

Und-PP-diNAcBac enzymatic synthesis (*Bacillus subtilis* EpsL) ................................ S14

**Figure S6.** *Bs* EpsL reaction after 30 minutes ............................................................. S14

**Table S3.** *B. subtilis* strains used in this study ............................................................ S16

Plasmids ........................................................................................................................ S17

**Table S4.** Plasmids used in the genetic section of this study ........................................ S18

**Table S5.** Primers used in the genetic section of this study .......................................... S18

**Table S6.** Custom synthesized gene sequences ........................................................... S19

**Figure S7.** Colony biofilm morphology and hydrophobicity upon genetically complementing *ΔepsL-Bs* mutant with *pglC* of *Campylobacter* ........................................................................ S23

**Table S7.** UniProt accession numbers and associated protein sequences of glycosyl transferases (GT) used for sequence identity ........................................................................................ S23

**Figure S8:** Colony biofilm morphology and hydrophobicity upon genetically complementing *ΔepsF-Bs ΔepsD-Bs* mutant with *pglA* of *Campylobacter* and *Neisseria* .............................. S25

**Figure S9:** Structural alignment of PglA of *C. jejuni* (fluorescent green), *Bs* EpsD (salmon pink) and *Bs* EpsF (dark pink) ........................................................................................ S26

Detergent solubilization of *Bs* EpsD ............................................................................ S27

**Figure S10.** SDS PAGE and Western blotting analysis of *Bs* EpsD cell envelope fraction ............ S27

**Figure S11.** *Bs* EpsD detergent solubilization with Triton X-100 visualized by SDS PAGE .......... S28

Und-PP-diNAcBac enzymatic synthesis (*Campylobacter concisus* PglC) ........................ S28

Radioactivity-based biochemical assays on *Bs* EpsD cell envelope fraction (CEF) .... S30

**Figure S12.** Substrate determination for the first glycosyl transferase in *B. subtilis,* EpsD, using a radioactivity-based assay ........................................................................................ S31

**Figure S13:** Comparison of *B. subtilis* EpsD with PglAs from *C. concisus* and *C. jejuni* ........ S32

References .................................................................................................................... S33

**General Information.** Unless otherwise specified, all commercially available reagents were used without further purification. Normal phase chromatographic purifications were conducted using SilicaFlash Irregular Silica Gel P60, 40 - 63 μm, 60Å (SiliCycle). Thin-layer chromatography (TLC) was performed on SilicaPlate glass-backed, silica gel TLC plates (250 μM, F254, SiliCycle TLG-R10014B-323). Some figures in this document were created with Biorender.com.

**Table S1.** UniProt accession numbers and associated protein sequences of monoPGTs used for sequence identity.

| | UniProt Accession number | Organism | Sequence | Protein (gene) |
|---|---|---|---|---|
| *Bs* EpsL | P71062 | *Bacillus subtilis* | MILKRLFDLTAAIFLLCCTSVIILFTIAVVRLKIGSPVFF KQVRPGLHGKPFTLYKFRTMTDERDSKGNLLPDEV RLTKTGRLIRKLSIDELPQLLNVLKGDLSLVGPRPLL MDYLPLYTEKQARRHEVKPGITGWAQINGRNAISW EKKFELDVWYVDNWSFFLDLKILCLTVRKVLVSEGIQ QTNHVTAERFTGSGDVSS | Uncharacterized sugar transferase EpsL (*epsL*) |
| *Hp* PglC | E1B268 | *Helicobacter pylori* | MYKNLIKPILDFILAFLLIIIFSPIILIVALLIKLKLGSPILFT QERPGLNGKIFRIYKFRTMSDERDSKGDLLSDELRL KGFGKLIRKSSLDELPQLFNVLKGEMSFVGPRPLLV EYLKLYNQEQAKRHNVKPGITGWAQVNGRNAISWE EKFKLDVYYVEHISFMLDCKILYMTFFKVLKRKDINS NTNITMEKFTGNKSE | Bacterial sugar transferase (*pglC*) |
| *Cj* PglC | Q0P9D0 | *Campylobacter jejuni* | MYEKVFKRIFDFILALVLLVLFSPVILITALLLKITQGSV IFTQNRPGLDEKIFKIYKFKTMSDERDEKGELLSDEL RLKAFGKIVRSLSLDELLQLFNVLKGDMSFVGPRPLL VEYLPLYNKEQKLRHKVRPGITGWAQVNGRNAISW QKKFELDVYYVKNISFLLDLKIMFLTALKVLKRSGVS KEGHVTTEKFNGKN | Undecaprenyl phosphate *N,N'*-diacetylbacillosamine 1-phosphate transferase (*pglC*) |
| *Cc* PglC | A7ZET4 | *Campylobacter concisus* | MYRNFLKRVIDILGALFLLILTSPIIIATAIFIYFKVSRDVI FTQARPGLNEKIFKIYKFKTMSDERDANGELLPDDQ RLGKFGKLIRSLSLDELPQLFNVLKGDMSFIGPRPLL VEYLPIYNETQKHRHDVRPGITGLAQVNGRNAISWE KKFEYDVYYAKNLSFMLDVKIALQTIEKVLKRSGVSK EGQATTEKFNGKN | *N,N'*-diacetylbacilliosaminyl-1-phosphate transferase (*pglC*) |

| | | | | |
|---|---|---|---|---|
| *Sa* Cap5M | P95706 | *Staphylococcus aureus* | MKRLFDVVSSIYGLVVLSPILLITALLIKMESPGPAIFKQKRPTINNELFNIYKFRSMKIDTPNVATDLMDSTSYITKTGKVIRKTSIDELPQLLNVLKGEMSIVGPRPALYNQYELIEKRTKANVHTIRPGVTGLAQVMGRDDITDDQKVAYDHYYLTHQSMMLDMYIIYKTIKNIVTSEGVHH | Bacterial sugar transferase (*cap5M*) |

Protein sequence of EpsL-$His_6$ from *Bacillus subtilis (Bs):*

MILKRLFDLTAAIFLLCCTSVIILFTIAVVRLKIGSPVFFKQVRPGLHGKPFTLYKFRTMTD
ERDSKGNLLPDEVRLTKTGRLIRKLSIDELPQLLNVLKGDLSLVGPRPLLMDYLPLYTEK
QARRHEVKPGITGWAQINGRNAISWEKKFELDVWYVDNWSFFLDLKILCLTVRKVLVS
EGIQQTNHVTAERFTGSGDVSSHHHHHH

**Overexpression of *Bs* EpsL and *Bs* EpsD.** EpsL and EpsD constructs were transformed into BL21(DE3)-RIL cells and overexpressed in the autoinduction medium. A 5 mL seed culture supplemented with 50 µg/mL kanamycin was grown in MDG medium at 37ºC for 18 hours at 225 rpm for each enzyme. The overnight seed cultures were inoculated in 500 mL autoinduction media (0.1% (w/v) tryptone, 0.05% (w/v) yeast extract, 2 mM $MgSO_4$, 0.05% (v/v) glycerol, 0.005% (w/v) glucose, 0.02% (w/v) $\alpha$-lactose, 2.5 mM $Na_2HPO_4$, 2.5 mM $KH_2PO_4$, 5 mM $NH_4Cl$, 0.5 mM $Na_2SO_4$) in a baffled flask, supplemented with 30 µg/mL chloramphenicol and 90 µg/mL kanamycin. The 500 mL cultures were incubated at 37 ºC for 4-5 hours at 225 rpm till the bacterial growth reached the log phase ($OD_{600}$ ~ 0.8-1). The incubation temperature was reduced to 16ºC for the autoinduction of the protein expression for 18 hours at 225 rpm. The cells were harvested at 3000 rpm for 25 minutes at 4ºC. The pellets were washed with 15 mL phosphate buffer saline, flash frozen in $LN_2$, and stored at –80ºC.

**Preparation of cell envelope fraction (CEF).** Cell pellets were resuspended in 50 mL 50 mM HEPES pH 7.5, 100 mM NaCl, with 25 mg lysozyme (Research Products International, cat # L38100), 25 µL DNAse I (New England BioLabs cat # M0303S), and 50 µL protease inhibitor cocktail (Roche, cat # 11836170001). Cells were sonicated twice for 1.5 minutes (1 second on/2 seconds off, 50% amplitude), resting on ice for 5 minutes in between sonication cycles. For EpsL, the lysed cells were centrifuged at 9000 rpm for 45 minutes (low-speed spin) using a 45-Ti rotor. The resulting supernatant was transferred to a clean centrifuge tube and centrifuged at 35,000 rpm for 65 minutes (high-speed spin) using a 45-Ti rotor to pellet the membrane fraction. For EpsD, the lysed cells were directly centrifuged at 35,000 rpm for 65 minutes (high-speed spin) using a 45-Ti rotor to pellet the membrane fraction. The CEF was homogenized (Dounce) into 12.5 mL 50 mM HEPES pH 7.5, 100 mM NaCl, with the addition of 14 µL of protease inhibitor cocktail.

**Detergent screening of *Bs* EpsL.** Small-scale detergent extraction of EpsL was conducted using Anatrace analytical extractor kit (part number AL-EXTRACT) according to the manufacturer's protocol with slight modifications. Each detergent at 5x stock solutions was diluted to the working 1x stocks in resuspension buffer (50 mM HEPES pH 7.5, 100 mM NaCl). The CEF (30 µL of 37 mg/mL *total protein*) of EpsL was diluted with each of the eight detergents (1x stocks, 150 µL) to a final volume of 180 µL. The CEF was solubilized at 4ºC for 2 h by a gentle rotation followed by centrifugation at 100,000 x g for 1 h using a Beckman-Coulter Ti 42.2 rotor and Beckman-Coulter open-top thick wall polypropylene tubes (7 x 20 mm, Part # 343621). The amount of solubilized protein was visualized by SDS PAGE and Western blotting analysis.

**Table S2.** The working concentration and structures of the detergents used in the solubilization screen.

| Detergent Name | CMC (% w/v) | Working concentration | Structure |
|---|---|---|---|
| Triton X-100 | 0.02% | 1% | 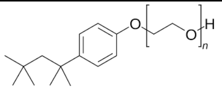 |
| $C_8E_4$ | 0.25% | 2% | 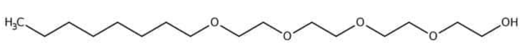 |
| $C_{12}E_8$ | 0.00% | 0.50% | 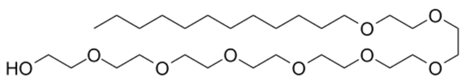 |
| DM | 0.09% | 1% | 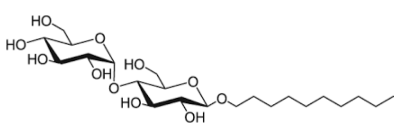 |

| | | | |
|---|---|---|---|
| DDM | 0.01% | 1% | 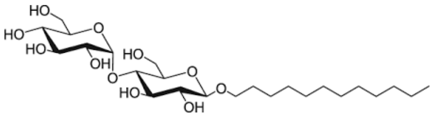 |
| FC-12 | 0.05% | 1% | 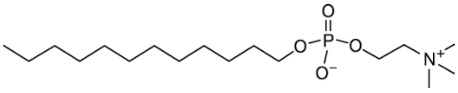 |
| LDAO | 0.02% | 1% | 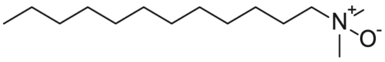 |
| OG | 0.53% | 2% | 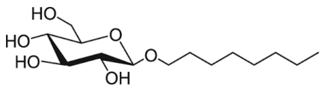 |

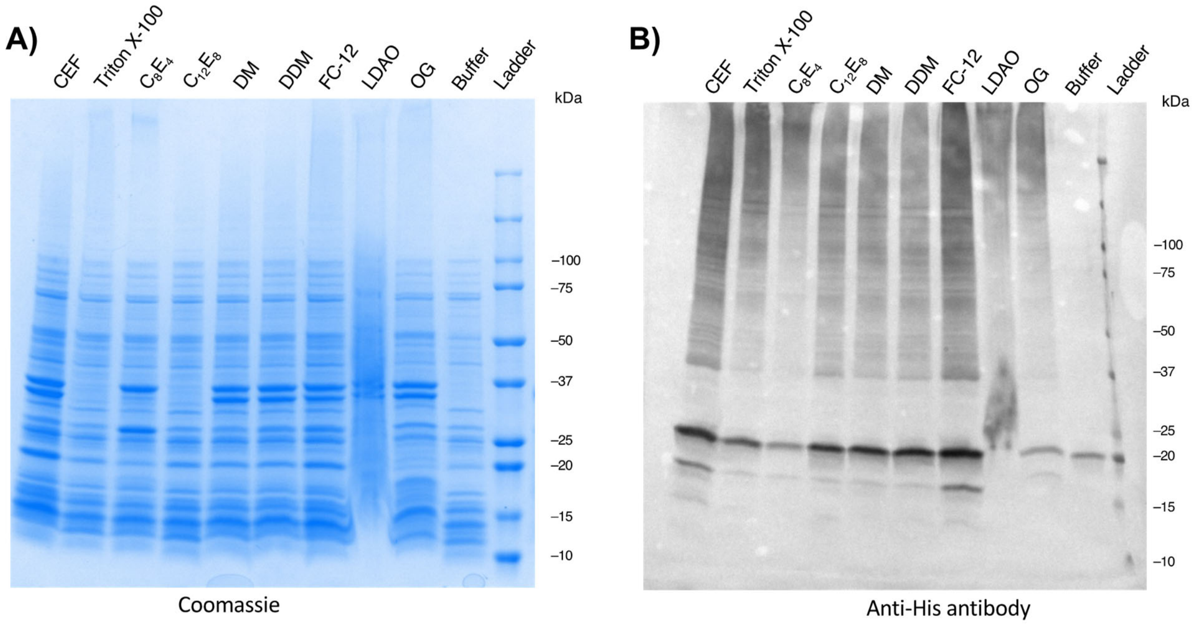


**Figure S1.** *Bs* EpsL detergent solubilization screen visualized by **A)** SDS-PAGE and **B)** Western blotting analysis.

**Purification of EpsL.** The CEF of EpsL in 5.5 mL of 50 mM HEPES pH 7.5, 100 mM NaCl, was solubilized into 0.09% Triton X-100 (Anapoe-X-100) or 0.03% Octaethylene Glycol Monododecyl Ether ($C_{12}E_8$) and incubated with rotation at 4ºC for 2 hours. The detergent-homogenized sample was centrifuged at 42,000 rpm for 65 minutes using a

70-Ti rotor. The supernatant was incubated with 1 mL Ni-NTA resin for 1 hour at 4ºC. The resin was washed with 10 mL Wash I buffer (50 mM HEPES pH 7.5, 100 mM NaCl, 15 mM imidazole, 5% glycerol, 0.09% Triton X-100 or 0.03% $C_{12}E_8$), followed by a wash with 10 mL Wash II buffer (50 mM HEPES pH 7.5, 100 mM NaCl, 45 mM imidazole, 5% glycerol, 0.09% TritonX-100 or 0.03% $C_{12}E_8$). EpsL was eluted in 2 x 0.5 mL fractions of elution buffer (50 mM HEPES pH 7.5, 100 mM NaCl, 500 mM imidazole, 5% glycerol, 0.09% TritonX-100 or 0.03% $C_{12}E_8$) and immediately desalted using a 5 mL desalting column in 50 mM HEPES pH 7.5, 100 mM NaCl, 5% glycerol, 0.09% TritonX-100 or 0.03% $C_{12}E_8$. Desalted protein was eluted in 0.5 mL fractions from 1.5 mL – 3.0 mL. Fractions were pooled, flash-frozen in $LN_2$, and stored at –80°C.

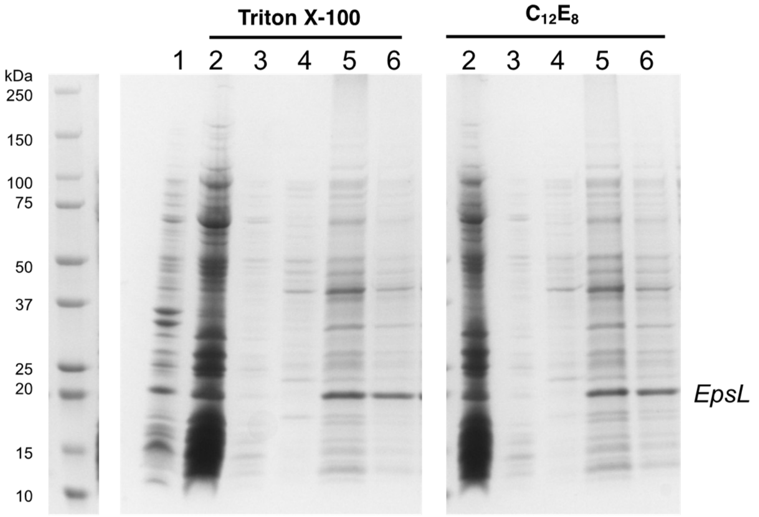


**Figure S2.** *Bs* EpsL detergent solubilization with Triton X-100 and $C_{12}E_8$ visualized by SDS PAGE (Coomassie). Lanes: 1) CEF; 2) solubilized protein; 3) flow through; 4) 45 mM imidazole wash; 5) 500 mM imidazole elution; 6) desalted pure fraction.

**General information for diphosphate sugars.** All commercially available uridine diphosphate sugars (UDP-sugars) were purchased from Millipore Sigma. Und-PP-diNAcBac was chemoenzymatically synthesized from UndP and UDP-diNAcBac (*vide infra)*. The [$^3$H]UDP-sugar substrates were diluted from the following specific activities (Ci/mmol) to provide a certain amount of disintegrations per minute (dpm) per assay: UDP-[$^3$H]Gal (40 Ci/mmol, 73000 dpm/assay), UDP-[$^3$H]Glc (60 Ci/mmol, 56000 dpm/assay), UDP-[$^3$H]GlcNAc (20 Ci/mmol, 67000 dpm/assay), UDP-[$^3$H]GalNAc (20

Ci/mmol, 65000 dpm/assay), and UDP-[$^3$H]diNAcBac (500 dpm/pmol, 67000 dpm/assay). Conversion of microcurie (µCi) to dpm is: 1 Ci = 2.22 x $10^{12}$ dpm.

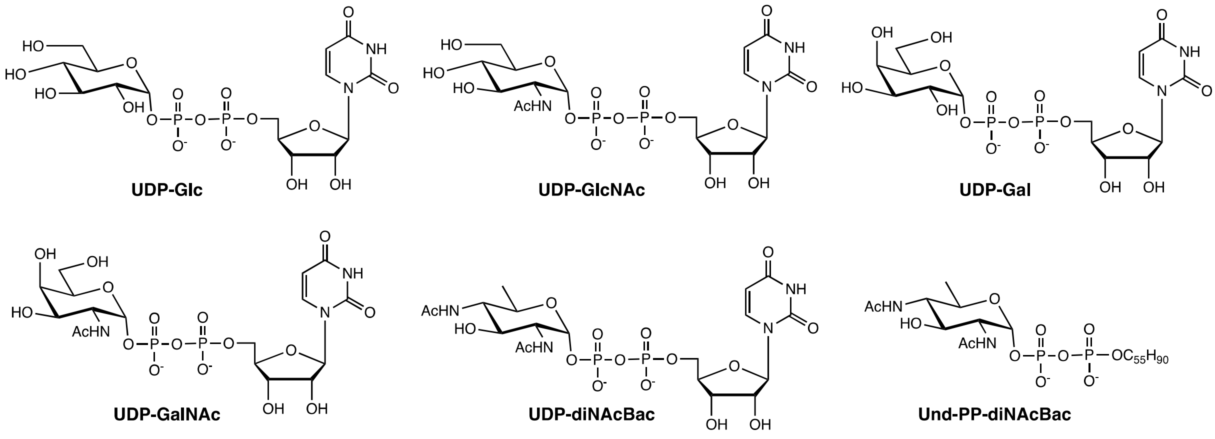


**Figure S3.** Uridine diphosphate sugars used in this study.

**Western blotting analysis.** Protein samples were separated by gel electrophoresis on Biorad 4−20% gradient gels. The samples were loaded for western blot and SDS-PAGE analyses. Western blot analyses were transferred at 100 V for 70 minutes at 4 °C. The membrane was then incubated in 25 mL of 3% BSA (0.75 g BSA in 25 mL TBS-T) for 30 minutes to prevent the nonspecific binding of antibodies to the membrane. For the detection of the $His_6$-tagged proteins, the membrane was incubated with a 1:50000 dilution of mouse anti-His antibody (LifeTein) in TBS-T 3% BSA (5 µL of 1mg/mL in 25 mL TBS-T with 3% BSA) for 1 hour. The membrane was washed with TBS-T for 5 minutes (5x) followed by incubation with a 1:10000 dilution of secondary goat anti-mouse antibody with alkaline phosphatase (AP) conjugate in TBS-T buffer (1.5 µL of 0.6 mg/mL in 15 mL TBS-T) for 1 hour. The solution was removed, and the membrane was washed with TBS-T (3 x 5 min), followed by TBS (3 x 5 min). The western blot was revealed with alkaline phosphatase substrate (1-step NBT/BCIP) and allowed to develop for 5 minutes. The blot was washed with water and imaged using a BioRad Molecular Imager Gel Doc XR+ (colorimetric).

**Protein expression for UDP-diNAcBac chemoenzymatic synthesis.**

**Expression of PglF $_{\Delta 1\text{-}130}$ from *C. jejuni* and PglC from *N. gonorrhoeae*.** PglF$_{\Delta 1\text{-}130}$ was transformed into BL21 cells and grown on an agar plate supplemented with 100 µg/mL carbenicillin. A 5 mL seed culture was grown from a single colony in LB medium, supplemented with 100 µg/mL carbenicillin at 37ºC for 18 hours at 225 rpm. The overnight seed cultures were inoculated in 500 mL MDG media (0.1% (w/v) tryptone, 0.05% (w/v) yeast extract, 2 mM $MgSO_4$, 0.05% (v/v) glycerol, 0.005% (w/v) glucose, 0.02% (w/v) α-lactose, 2.5 mM $Na_2HPO_4$, 2.5 mM $KH_2PO_4$, 5 mM $NH_4Cl$, 0.5 mM $Na_2SO_4$) in a baffled flask, supplemented with 100 µg/mL carbenicillin. The 500 mL cultures were incubated at 37ºC for 4-5 hours at 225 rpm until the bacterial growth reached the log phase ($OD_{600}$ ~ 0.8-1). The incubation temperature was reduced to 16ºC and the protein expression was induced upon the addition of 1 mM IPTG. The culture was incubated for 18 hours at 225 rpm for protein expression. PglC (*Ng*) was transformed into BL21(DE3) cells and grown on an agar plate supplemented with 50 µg/mL kanamycin. A 5 mL seed culture was grown from a single colony in LB medium, supplemented with 50 µg/mL kanamycin at 37ºC for 18 hours at 225 rpm. The overnight seed cultures were inoculated in 500 mL MDG media (0.1% (w/v) tryptone, 0.05% (w/v) yeast extract, 2 mM $MgSO_4$, 0.05% (v/v) glycerol, 0.005% (w/v) glucose, 0.02% (w/v) α-lactose, 2.5 mM $Na_2HPO_4$, 2.5 mM $KH_2PO_4$, 5 mM $NH_4Cl$, 0.5 mM $Na_2SO_4$) in a baffled flask, supplemented with 50 µg/mL kanamycin. The 500 mL cultures were incubated at 37ºC for 4-5 hours at 225 rpm until the bacterial growth reached the log phase ($OD_{600}$ ~ 0.8-1). The incubation temperature was reduced to 16ºC for the autoinduction of the protein expression for 18 hours at 225 rpm. The cells were harvested at 3000 rpm for 25 minutes at 4ºC. The pellets were washed with 15 mL phosphate buffer saline, flash-frozen in $LN_2$, and stored at –80ºC.

**Immobilization of PglF $_{\Delta 1\text{-}130}$ on Glutathione-resin.** BL21 cells from 0.5 L cultures with overexpressed GST- PglF $_{\Delta 1\text{-}130}$ were thawed on ice and resuspended in 40 mL lysis buffer (50 mM HEPES, pH 7.5, 150 mM NaCl, 25 µg lysozyme, 25 µL DNAse I, 40 µL protease inhibitor cocktail) by incubating at 4ºC for 30 minutes with gentle rotation. Cells were sonicated for 90 seconds with 50 % amplitude with 1 s ON and 2 s OFF cycles. The homogenized lysate was transferred into an ultracentrifuge tube and centrifuged at 35,000 rpm in a Ti45 rotor for 1 h at 4ºC. The clarified lysate was transferred into a clean tube and incubated with 4 mL Glutathione agarose resin (Pierce™), pre-equilibrated with working buffer (50 mM HEPES, pH 7.5, 150 mM NaCl). The resin and lysate mixtures were incubated with $NAD^+$ at a final concentration of 1 mM for 4 h at 4°C. The resin was transferred into a chromatographic column and the excess clarified lysate was flowed through the column via gravity. The column was washed with 8 column volumes (CV) of working buffer at 4ºC to remove excess protein and the immobilized GST- PglF $_{\Delta 1\text{-}130}$ was used immediately.

**Immobilization of PglC on Ni-NTA resin.** BL21(DE3) cells from 0.5 L cultures with overexpressed PglC-$His_6$ were thawed on ice and resuspended in 40 mL lysis buffer (50 mM HEPES, pH 7.5, 150 mM NaCl, 25 µg lysozyme, 25 µL DNAse I, 40 µL protease inhibitor cocktail) by incubating at 4ºC for 30 minutes with gentle rotation. Cells were sonicated for 90 s with 50 % amplitude with 1 s ON and 2 s OFF cycles. The homogenized lysate was transferred into an ultracentrifuge tube and centrifuged at 35,000 rpm in a Ti45 rotor for 1 h at 4ºC. The clarified lysate was transferred into a clean tube and incubated with 4 mL Ni-NTA agarose resin (Pierce™), pre-equilibrated with working buffer (50 mM HEPES, pH 7.5, 150 mM NaCl). The resin was incubated for 4 h at 4ºC with pyridoxal phosphate (PLP) at a final concentration of 1 mM. The resin was transferred into a chromatographic column and the excess clarified lysate was flowed through the column via gravity. The column was washed with 8 CV of working buffer at 4ºC with 15 mM imidazole, followed by 2 CV of working buffer at 4ºC without imidazole to remove excess protein. The immobilized PglC-$His_6$ was used immediately.

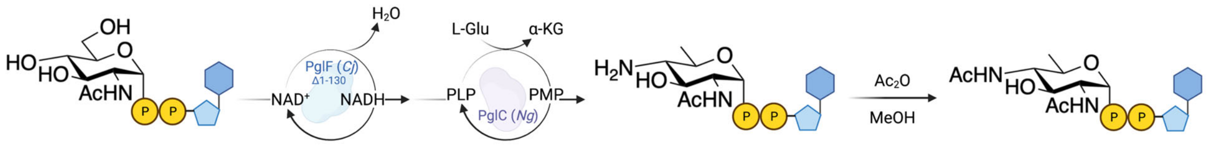


**Scheme S1.** Chemoenzymatic synthesis of UDP-diNAcBac.

**UDP-diNAcBac chemoenzymatic synthesis.** Glutathione agarose resin with immobilized GST-PglF$_{\Delta 1\text{-}130}$ and Ni-NTA agarose resin with immobilized PglC-$His_6$ was resuspended in 1 CV of reaction buffer (50 mM HEPES, pH 8, 150 mM NaCl) and combined in a 50 mL conical tube. UDP-GlcNAc (25 mg) was dissolved in the reaction buffer and added to the resin mix. Additional $NAD^+$ and PLP were added to the resin mix at a final concentration of 0.5 mM each. Additionally, L-glutamic acid was added at a final concentration of 25 nM and the reaction proceeded for 68-70 h at room temperature with gentle rotation. The resin mix was transferred to a chromatographic column and the product was collected in the flow-through and combined with the resin washes (2 CV of reaction buffer). Protein present in the flow-through and wash fractions were removed by heating the solution at 60ºC for 1 h, followed by centrifugation at 3,200 x g for 30 minutes. The crude UDP-4-aminosugar was purified using a Waters Sep-Pak C18 3cc Vac Cartridge (Silica-based, 200 mg Sorbent, 55-105 µm, Waters Corp WAT054945). The compound was loaded and eluted in $H_2O$ (0.1% TFA) and visualized by TLC (UV 254 nm, mobile phase: (5:1:3:1) *n*-BuOH/EtOAc/$H_2O$/25% ammonium hydroxide). The combined fractions were lyophilized to provide 22.7 mg of UDP-4-amino sugar (93% yield). The yield was determined by UV-VIS at 262 nm with the extinction coefficient of 10,000 $M^{-1}$ $cm^{-1}$. A fraction of the UDP-4-aminosugar stock (2.11 mg) was then dissolved in MeOH (0.5 mM of UDP-sugar in MeOH), followed by the addition of $Ac_2O$ (40 equivalents) and rotated at ambient temperature for 3 h. The chemical acetylation was monitored by TLC (mobile phase: (5:1:3:1) *n*-BuOH/EtOAc/$H_2O$/25% ammonium hydroxide, UV 254 nm) and after complete consumption of starting material, the reaction mixture was concentrated in volume under a stream of $N_2$. The resulting crude mixture was purified using a Waters Sep-Pak C18 3cc Vac Cartridge and eluted in $H_2O$ (0.1% TFA). The combined fractions containing the desired product were lyophilized to yield UDP-diNAcBac in 72% yield (1.64 mg).

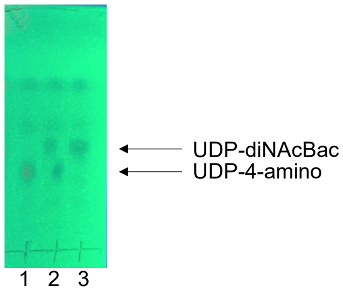


**Figure S4.** Thin-layer chromatography (TLC) of the chemical acetylation reaction of UDP-4-amino sugar. The solvent system used was (5:1:3:1) *n*-BuOH/EtOAc/$H_2O$/25% ammonium hydroxide and visualized by UV (254 nm). Lanes: 1) UDP-4-amino sugar starting material, 2) Co-spot, and 3) reaction after 3 h.

**UDP-[$^3$H]diNAcBac enzymatic synthesis and purification.** UDP-[$^3$H]diNAcBac was synthesized using a modified literature procedure (1, 2). To synthesize UDP-[$^3$H]diNAcBac, 200 nmol of purified UDP-4-aminosugar was incubated with 2.5 nmol [$^3$H]AcCoA (20 Ci/mmol, American Radiolabeled Chemicals) and 26 µM PglD in 50 mM HEPES, pH 7.5, 100 mM NaCl for 30 minutes at room temperature, followed by a chase with an excess of non-radiolabeled AcCoA (247.5 nmol). After overnight rotation, the reaction was supplemented with an additional 13 µM PglD and was allowed to proceed for two more hours. The reaction mixture was prepared for HPLC purification by heating at 60˚C for 1 hour and centrifugation at 16,000 x g for 10 minutes to precipitate and remove protein. The radiolabeled product was purified on a semi-preparative Dionex CarboPac PA1 HPLC column using the following method: Buffer A = water; Buffer B = 1 M $NH_4HCO_3$; 10%-20% B over 20 minutes, 20%-70% B over 1 minute, 70% B for 10 minutes, 70%-10% B over 1 minute, 10% B for 15 minutes. The fractions containing UDP-[$^3$H]diNAcBac were pooled and lyophilized for several rounds to remove the $NH_4HCO_3$.

**Radioactivity-based biochemical assays on purified protein (*Bs* EpsL)**

EpsL *(Bs)* substrate specificity was measured using a radioactively-labeled UDP-[$^3$H]-sugar panel and an extraction-based assay (3). Enzymatic reactions contained 20 µM UndP (2.5 µL of 200 µM in DMSO), UDP-[$^3$H]-sugar (1 µL in $H_2O$), and 4.48 µM detergent-solubilized EpsL (2 µL of 56 µM) in a final volume of 25 µL of assay buffer (19.5 µL of 50 mM HEPES pH 7.5, 100 mM NaCl, 0.1% Triton X-100, 5 mM $MgCl_2$). The assay contained a final concentration of 10% DMSO. UndP and UDP-[$^3$H]-sugar were pre-incubated in the assay buffer for 30 seconds and the reactions were initiated by the addition of EpsL (*Bs*). An aliquot of 15 µL was quenched into 1 mL of 2:1 $CHCl_3$/MeOH after 15 minutes. The organic layer was washed three times with 500 µL PSUP (Pure Solvent Upper Phase = 15 mL $CHCl_3$, 240 mL MeOH, 1.83 g KCl, 235 mL $H_2O$). The organic layer was mixed with 5 mL Opti-Fluor O scintillation cocktail (PerkinElmer) and the combined aqueous layers were mixed with 5 mL EcoLite Liquid Scintillation Cocktail (MP Biomedicals). All layers were analyzed on a Beckman Coulter LS6500 scintillation counting system with quench compensation. EpsL *(Bs)* activity is background subtracted and reported as both dpm in the organic layer, and the percentage of dpm in the organic layer normalized to the total dpm per quenched point. Error bars represent biological duplicates and were calculated with GraphPad Prism 8.

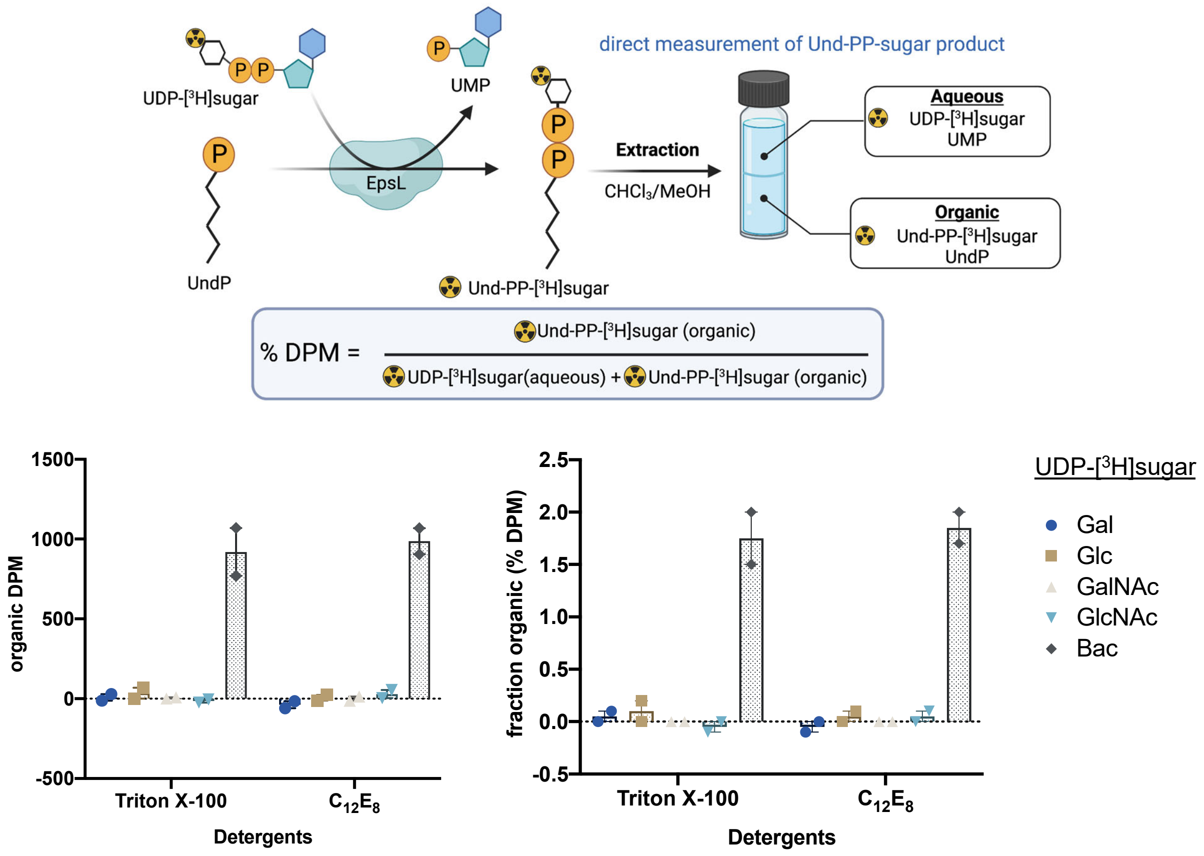


**Figure S5**. Substrate specificity determination with *B. subtilis* EpsL using a radioactivity-based assay. Error bars are given for mean ± SEM, n = 2.

**UMP Glo biochemical assays.** *B. subtilis* EpsL assays were performed using the Promega UMP-Glo assay, which detects UMP generated over the course of the reaction. The quenching solution was prepared as described by Promega. A UMP-Glo standard curve was obtained using final [UMP] concentrations of 10 μM, 5 μM, 2.5 μM, 1.25 μM, 0.625 μM, 0.3125 μM, 0.15625 μM, and 0 μM from 10x UMP stocks. The standard curve contained 10% DMSO. The EpsL assays contained 5.6 μM EpsL, 20 μM UndP (10% DMSO final), 0.1% Triton X-100, 50 mM HEPES at pH 7.5, 100 mM NaCl, 5 mM $MgCl_2$, and 50 μM UDP-sugar in a final volume of 11 μL. EpsL was preincubated in the reaction mixture lacking the UDP-sugar for 5 minutes at ambient temperature. Upon the addition of the UDP-sugar, the reaction was allowed to proceed for 30 minutes before the addition of the quenching solution. The reaction mixture was transferred to a 96-well plate (white, nonbinding surface, Corning). The plate was shaken at low speed for 30 s and incubated for 1 h at 25 ºC, and luminescence was read on the plate reader. Error bars represent biological triplicate and were calculated with GraphPad Prism 8.

**Und-PP-diNAcBac enzymatic synthesis (*Bacillus subtilis* EpsL).** The Und-PP-Bac reaction was set up in a 7 mL scintillation vial. The reaction contained a total volume of 200 µL and consisted of 25 µM UndP, 50 µM UDP-diNAcBac, 5.6 µM *Bs* EpsL, 50 mM HEPES pH 7.5, 100 mM NaCl, 0.1% Triton X-100 and 5 mM $MgCl_2$. The reactions contained a final concentration of 10% DMSO. The reaction was allowed to proceed for 30 minutes and monitored by UMP Glo and thin-layer chromatography (TLC).

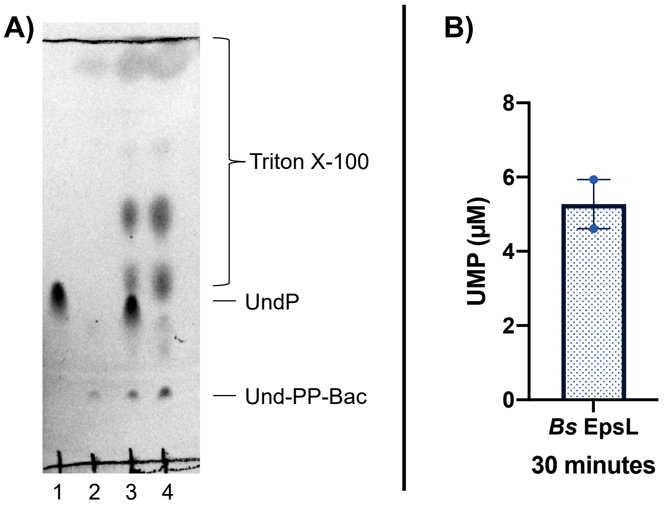


**Figure S6.** *Bs* EpsL reaction after 30 minutes. **A)** Thin-layer chromatography (TLC) with a solvent system of (65:25:4) $CHCl_3$/MeOH/$H_2O$. Lanes: 1) UndP standard, 2) Und-PP-Bac standard produced from *Cc* PglC, 3) Co-spot, and 4) Und-PP-Bac reaction with *Bs* EpsL. TLC is visualized by CAM staining and imaged on a BioRad Molecular Imager Gel Doc XR+. *Note: The reaction conversion cannot be quantified by TLC due to the differences in staining capabilities between UndP and Und-PP-Bac. Representation of conversion is more accurately visualized by UMP Glo. **B)** UMP Glo activity assay of the EpsL Und-PP-Bac reaction after 30 minutes. Error bars are given for mean ± SEM, n = 2.

**Bacterial strains and growth conditions:** All *B. subtilis* strains used and constructed in this study are listed in **Table S3**. *E. coli* and *B. subtilis* strains were routinely grown in LB medium (10 g NaCl, 5 g yeast extract, and 10 g tryptone per liter). Complex colony biofilms were grown on MSgg agar medium (5 mM potassium phosphate and 100 mM

MOPs at pH 7.0 supplemented with a metal mix containing 2 mM $MgCl_2$, 0.7 mM $CaCl_2$, 50 µM $MnCl_2$, 50 µM $FeCl_3$,1 µM $ZnCl_2$, 2 µM thiamine, 0.5% (v/v) glycerol, and 0.5% (w/v) glutamic acid solidified with 1.5% (w/v) Select Agar (Invitrogen)) (4, 5). The biofilms were grown at 30 °C for 48 h. Ectopic gene expression was induced with 25 µM isopropyl β-D-1-thiogalactopyranoside (IPTG). When appropriate, the antibiotics were used at the following concentrations: ampicillin: 100 µg/ml and spectinomycin:100 µg/ml.

**Strain Construction:** All strains, plasmids, and primers used in this study are presented in **Table S3**, **Table S4** and **Table S5**. *E. coli* strain MC1061 [*F'lacIQ lacZM15 Tn10 (tet)*] was used for the construction and maintenance of all the plasmids. The custom synthesized genes *pglC*$^{Cc}$, *pglC*$^{Cj}$, *pglA*$^{Cj}$, *pglA*$^{Ng}$, and *pglA*$^{Nm}$ were codon optimized for optimum expression in *B. subtilis* and were cloned in pUC57 standard plasmid by Genscript using SalI and SphI restriction sites. **Table S6** provides more details of the sequences synthesized. The plasmids received from Genscript were used to digest the synthesized gene and cloned into pDR111 plasmid using SalI and SphI restriction sites to generate plasmids pNW2127, pNW1931, pNW1923, pNW1932 and pNW1933 respectively. The *epsL, epsF* and *epsD* coding sequences of *B. subtilis* were also cloned into pDR111 to generate pNW2100, pNW2109, and pNW2103 plasmids. These plasmids were introduced into *B. subtilis* 168 genome using competent cells generated with standard protocols (6). The plasmids integrated into *B. subtilis* chromosome at the non-essential *amyE* gene locus and the coding region was placed under the control of IPTG-inducible promoter, Phy-spank. SPP1 phage preparation and transduction to introduce DNA into *B. subtilis* strain NCIB 3610 were conducted as described previously (7).

**Colony biofilm morphology assay:** *B. subtilis* strains were streaked on LB agar plates and incubated overnight at 37 °C. The following day single colonies were grown in 3 ml of LB broth at 37 °C with agitation until an $OD_{600}$ ≈1.0. All the cultures were normalized to the same density and 5 µl of the cultures were spotted onto MSgg media plates, without and with 25 µM IPTG. The plates were incubated at 30 °C for 48 h before imaging. For all the strains, three independent biological replicates along with their two technical replicates were set up. Biofilm imaging was performed using an MZ16 FA stereomicroscope (Leica) using LAS version 2.7.1. The images were imported into the OMERO server for data management and analysis (8).

**Quantification of biofilm surface area:** To quantify the surface area or footprints of biofilms Fiji/ImageJ software (9, 10) was used with a recently established macro (11, 12) that uses built-in function of ImageJ to detect biofilm regions. The images of colony biofilms were saved as multi-series Leica .LIF files after stereoscopic imaging. The .LIF file was uploaded to macro in Fiji to import the data and the batch analysis was done on the brightfield images. The outcome was a summary table of detected surface area of biofilms above the background. A minimum of three biological and two technical replicates were performed for each strain.

**Biofilm hydrophobicity assay:** The hydrophobicity of biofilms was tested by measuring the contact angle between the surface of the biofilm grown at 30 °C for 48 h and a 5 µl water drop of water, as described previously (13). The measurements were taken 5 minutes after the initial placement of the water droplet on the biofilm surface using a ThetaLite TL100 optical tensiometer (Biolin Scientific). The measurements were taken at 0 minutes in case of the absence of biofilm. Contact angles were determined with OneAttension software, using the Young-Laplace equation. Contact angles above 90° are indicative of a hydrophobic surface, whereas contact angles below 90° are considered hydrophilic. A minimum of three biological and two technical replicates were performed for each strain.

**Table S3: *B. subtilis* strains used in this study**

| **Strain number** | **Genotype**[a] | **Strain Construction/Reference**[b] |
|---|---|---|
| NCIB 3610 | Prototroph | B.G.S.C. |
| 168 | *trpC2* | B.G.S.C. |
| NRS5904 | NCIB 3610 Δ*epsF* | Ref(14) |
| NRS5905 | NCIB 3610 Δ*epsD* | Ref(14) |
| NRS5907 | NCIB 3610 Δ*epsL* | Ref(14) |
| NRS5930 | NCIB 3610 Δ*epsD amyE-Phy-spank-epsD-lacI* (spc) | SSP1 NRS5928 → NRS5905 |
| NRS5942 | NCIB 3610 Δ*epsL amyE-Phy-spank-epsL- lacI* (spc) | SSP1 NRS5941 → NRS5907 |
| NRS5961 | NCIB 3610 Δ*epsF amyE-Phy-spank-epsF- lacI* (spc) | SSP1 NRS5957→ NRS5904 |
| NRS5992 | NCIB 3610 Δ*epsL amyE-Phy-spank-pglC*$^{Cc}$*-lacI* (spc) | SSP1 NRS5990 → NRS5907 |

| | | |
|---|---|---|
| NRS6618 | NCIB 3610 Δ*epsL amyE-Phy-spank-pglC*$^{Cj}$*-lacI* (spc) | SSP1 NRS6613 → NRS5907 |
| NRS6605 | NCIB 3610 Δ*epsD amyE-Phy-spank-pglA*$^{Cj}$*-lacI* (spc) | SSP1 NRS6602 → NRS5905 |
| NRS6619 | NCIB 3610 Δ*epsD amyE-Phy-spank-pglA*$^{Ng}$*-lacI* (spc) | SSP1 NRS6614→ NRS5905 |
| NRS6620 | NCIB 3610 Δ*epsD amyE-Phy-spank-pglA*$^{Nm}$*-lacI* (spc) | SSP1 NRS6615 → NRS5905 |
| NRS6628 | NCIB 3610 Δ*epsF amyE-Phy-spank-pglA*$^{Cj}$*-lacI* (spc) | SSP1 NRS6602 → NRS5904 |
| NRS6629 | NCIB 3610 Δ*epsF amyE-Phy-spank-pglA*$^{Ng}$*-lacI* (spc) | SSP1 NRS6614 → NRS5904 |
| NRS6630 | NCIB 3610 Δ*epsF amyE-Phy-spank-pglA*$^{Nm}$*-lacI* (spc) | SSP1 NRS6615 → NRS5904 |
| NRS5928 | 168 *trpC2 amyE-Phy-spank-epsD-lacI* (spc) | pNW2100 → 168 |
| NRS5941 | 168 *trpC2 amyE-Phy-spank-epsL-lacI* (spc) | pNW2103 → 168 |
| NRS5957 | 168 *trpC2 amyE-Phy-spank-epsF-lacI* (spc) | pNW2109 → 168 |
| NRS5990 | 168 *trpC2 amyE-Phy-spank-pglC*$^{Cc}$*-lacI* (spc) | pNW2127 → 168 |
| NRS6613 | 168 *trpC2 amyE-Phy-spank-pglC*$^{Cj}$*-lacI* (spc) | pNW1931 → 168 |
| NRS6602 | 168 *trpC2 amyE-Phy-spank-pglA*$^{Cj}$*-lacI* (spc) | pNW1923 → 168 |
| NRS6614 | 168 *trpC2 amyE-Phy-spank-pglA*$^{Ng}$*-lacI* (spc) | pNW1932 → 168 |
| NRS6615 | 168 *trpC2 amyE-Phy-spank-pglA*$^{Nm}$*-lacI* (spc) | pNW1933 → 168 |

[a] The relevant genotypes are provided here. Spc, spectinomycin resistance cassette.
[b] The direction of strain construction is indicated with plasmid DNA or phage (SPP1) (→) recipient using standard techniques. Table 2 provides the details of the plasmids used here. B.S.G.C. represents the Bacillus genetic stock center.

**Plasmids.** The PglF gene without the N-terminal transmembrane domains (aa 1-130) from *Campylobacter jejuni* 11168 was cloned in pGEX-4T-2 vector with an N-terminal GST tag. The aminotransferase domain of PglC from *Neisseria gonorrhoeae* was cloned with a C-terminal 6x Histidine tag in vector pET30b(+). EpsD construct was purchased from Twist Bioscience with a C-terminal 6x histidine tag in pET24a vector between BamHI and HindIII sites. The EpsL construct was cloned into pET24a vector with a C-terminal 6x histidine tag using the Gibson assembly method. Genes that encode for *Cj* PglF, *Ng* PglC, *Bs* EpsL, and *Bs* EpsD were codon optimized for *E. coli*.

The primers used were:

5'-
GTTTAACTTTAAGAAGGAGATATACATATGATCCTCAAACGGCTGTTCGATCTTACT
GCGGCAATC-3' (forward) and

5'-GTGGTGGTGGTGGTGCGACGAAACGTCACC-3' (reverse).

**Table S4: Plasmids used in the genetic section of this study.**

| Plasmid number | Relevant Details |
|---|---|
| pNW2100 | pDR111-*epsD* |
| pNW2103 | pDR111-*epsL* |
| pNW2109 | pDR111-*epsF* |
| pNW2127 | pDR111-*pglC*$^{Cc\ (*)}$ |
| pNW1931 | pDR111-*pglC*$^{Cj\ (*)}$ |
| pNW1923 | pDR111-*pglA*$^{Cj\ (*)}$ |
| pNW1932 | pDR111-*pglA*$^{Ng\ (*)}$ |
| pNW1933 | pDR111-*pglA*$^{Nm\ (*)}$ |

(*) Genetic material was synthesized by GenScript. Refer to **Table S6** in Supplemental information for full details.

**Table S5: Primers used in the genetic section of this study.**

| Primer Name | Sequence 5'-3' [a] | Use |
|---|---|---|
| NSW2592 | GCAGCGTCATTCCAATTTTCAAAAAACAG | *epsD* deletion check |
| NSW2593 | GTGCGGTAGGCACTCGCTCTCA | |
| NSW2596 | AAATAAAAACCTGCCCGCATCCT | *epsF* deletion check |
| NSW2597 | TTTAGAAAAGCACCGACACTATCAGGTGG | |
| NSW2608 | GCACCGCTTTTATTTATTCATGCCG | *epsL* deletion check |
| NSW2609 | TTCCGACGTGAGCGCCTTCCT | |
| NSW872 | AGGTGTGGCATAATGTGTGTAATTGTGAGC | *amyE* pDR111 sequencing |
| NSW873 | TGAACAATCACGAAACAATAATTGGTACGTACG | |
| NSW2636 | CGGGTCGACTTAGGGGGAGTGAAAATGACGAAAAAGATATTGT | *epsD* cloning |
| NSW2637 | GCGGCATGCTCATACGCTTTTCTCCTTTGTATCCATATCCATGTAC | |
| NSW2642 | GGCGTCGACACGAAAGGAGCTGTGAATCTTTGATCC | *epsL* cloning |
| NSW2629 | CCGGCATGCTCATGAGGACACATCTCCGCTT | |
| NSW2653 | CGCGCATGCGAATTCGGAAGGGCTTGTCAAGCATGAATAGC | *epsF* cloning |
| NSW2654 | GGCGCATGCCGGAAAAGGACCATAACCGATGA | |

[a] The underlined sequence represents the restriction enzyme site used for cloning.

### Table S6: Custom synthesized gene sequences.

| Gene name | Strain Name | Restriction sites added for cloning | Plasmid generated | Original sequence, before codon optimization<br>Red: Restriction sites<br>Green: Ribosomal binding site<br>Black: codon optimized for *B. subtilis* | Codon optimized sequence, received from Genscript<br>Red: Restriction sites<br>Green: Ribosomal binding site<br>Black: codon optimized for *B. subtilis* |
|---|---|---|---|---|---|
| $pglC^{Cc}$ | *Campylobacter concisus* GenBank: QPI05301.1 | 5'-SalI-$pglC^{Cc}$-SphI-3' | pNW2127 | GTCGACAAGGAGGTGATCATT<br>AAAAATGTATAGAAATTTTTTA<br>AAGAGAGTGATTGATATTTTG<br>GGAGCTTTGTTTTTGCTCATTT<br>TAACATCGCCTATCATCATAG<br>CAACGGCGATTTTTATCTATTT<br>TAAGGTTAGCCGTGATGTCAT<br>TTTTACGCAGGCAAGGCCAGG<br>GCTAAATGAGAAAATTTTTAAA<br>ATTTATAAATTTAAGACGATGA<br>GCGACGAGCGTGACGCAAAT<br>GGCGAGCTCTTGCCAGATGAT<br>CAGCGTCTTGGTAAATTTGGC<br>AAACTTATCCGCTCACTTAGC<br>CTCGATGAGCTGCCACAGCTA<br>TTTAACGTGCTAAAGGGCGAT<br>ATGAGTTTCATCGGACCAAGG<br>CCGCTTTTGGTCGAGTACCTA<br>CCCATCTATAACGAAACGCAA<br>AAGCACCGCCACGACGTGCG<br>CCCTGGTATCACGGGTCTAGC<br>GCAGGTAAATGGCAGAAACG<br>CCATAAGCTGGGAGAAAAAAT<br>TTGAGTACGACGTCTATTATG<br>CTAAAAATTTAAGCTTTATGCT<br>TGATGTAAAGATCGCTTTGCA<br>GACCATCGAAAAAGTGCTAAA<br>ACGAAGTGGTGTCAGCAAAGA<br>GGGGCAGGCGACGACGGAGA<br>AATTTAATGGCAAAAACTAAG<br>CATGC | GTCGACAAGGAGGTGATCATTAAAAATGTACCGTA<br>ACTTCCTTAAACGTGTTATCGATATCCTTGGCGCTC<br>TTTTCCTTCTTATCCTTACATCTCCTATCATCATCGC<br>TACAGCTATCTTCATCTACTTCAAAGTTTCTCGTGA<br>TGTTATCTTCACACAAGCTCGTCCTGGCCTTAACGA<br>AAAAATCTTCAAAATCTACAAATTCAAAACAATGTCT<br>GATGAACGTGATGCTAACGGCGAACTTCTTCCTGA<br>TGATCAACGTCTTGGCAAATTCGGCAAACTTATCC<br>GTTCTCTTTCTCTTGATGAACTTCCTCAACTTTTCAA<br>CGTTCTTAAAGGCGATATGTCTTTCATCGGCCCTC<br>GTCCTCTTCTTGTTGAATACCTTCCTATCTACAACG<br>AAACACAAAAACATCGTCATGATGTTCGTCCTGGC<br>ATCACAGGCCTTGCTCAAGTTAACGGCCGTAACGC<br>TATCTCTTGGGAAAAAAAATTCGAATACGATGTTTA<br>CTACGCTAAAAACCTTTCTTTCATGCTTGATGTTAA<br>AATCGCTCTTCAAACAATCGAAAAAGTTCTTAAACG<br>TTCTGGCGTTTCTAAAGAAGGCCAAGCTACAACAG<br>AAAAATTCAACGGCAAAAACTAAGCATGC |
| $pglC^{Cj}$ | *Campylobacter jejuni* subsp. jejuni 81-176 GenBank: AAD51385.1 | 5'-SalI-$pglC^{Cj}$-SphI-3' | pNW1931 | GTCGACAAGGAGGTGATCATT<br>AAAAATGTATGAAAAAGTTTTT<br>AAAAGAATTTTTGATTTTATTTT<br>AGCTTTAGTGCTTTTAGTACTT<br>TTTTCTCCGGTGATTTTAATCA<br>CTGCTTTACTTTTAAAAATCAC<br>TCAAGGAAGTGTGATTTTCAC<br>TCAAAATCGCCCTGGGTTAGA<br>TGAAAAAATTTTTAAAATTTAT<br>AAATTTAAAACCATGAGCGAT<br>GAAAGAGATGAGAAGGGTGA<br>GTTATTAAGCGATGAATTGCG<br>TTTGAAAGCCTTTGGAAAAATT<br>GTTAGAAGCTTAAGTTTGGAT<br>GAGCTTTTGCAACTTTTTAATG<br>TTTTAAAAGGGGATATGAGTTT<br>TGTGGGGCCTAGGCCTCTTTT<br>GGTTGAGTATTTATCCCTTTAT<br>AATGAAGAGCAAAAATTGCGC<br>CATAAGGTGCGTCCAGGTATA<br>ACAGGATGGGCGCAGGTAAA<br>TGGCAGAAATGCTATTTCTTG<br>GCAGAAAAAATTCGAACTTGA<br>TGTGTATTATGTGAAAAATATT<br>TCTTTTTTGCTTGATTTAAAAA<br>TCATGTTTTTAACAGCTTTAAA<br>GGTTTTAAAACGAAGCGGGGT<br>AAGCAAAGAAGGCCATGTTAC | GTCGACAAGGAGGTGATCATTAAAAATGTATGAAA<br>AAGTTTTTAAAAGAATCTTTGATTTTATCCTTGCTTT<br>AGTTCTGCTTGTGTTATTTTCTCCGGTGATTCTGAT<br>CACAGCCTTACTGCTTAAAATTACACAAGGCTCAGT<br>CATCTTTACACAGAATAGACCGGGACTGGATGAAA<br>AAATCTTTAAAATCTACAAATTTAAAACAATGAGCG<br>ATGAACGCGATGAAAAAGGCGAATTACTGTCTGAT<br>GAACTGAGACTTAAAGCATTTGGAAAAATTGTCCG<br>CTCATTAAGCCTGGATGAACTTCTGCAACTGTTTAA<br>CGTTCTTAAAGGCGATATGTCATTTGTGGGACCGA<br>GACCGCTGCTTGTCGAATACCTTAGCCTGTACAAC<br>GAAGAACAGAAACTTAGACATAAAGTTCGCCCGGG<br>CATTACAGGATGGGCACAAGTGAATGGCCGCAAC<br>GCGATCAGCTGGCAGAAAAAATTTGAACTTGATGT<br>CTACTACGTTAAAAACATTTCATTTCTGCTGGATCT<br>GAAAATCATGTTTCTGACAGCTCTTAAAGTGTTAAA<br>ACGCTCTGGCGTCTCAAAAGAAGGACATGTTACAA<br>CAGAAAAATTTAATGGAAAAAACTAAGCATGC |

| | | | | | |
|---|---|---|---|---|---|
| | | | | AACAGAGAAATTTAATGGCAA<br>GAACTGAGCATGC | |
| *pglA*$^{Cj}$ | *Campylo bacter jejuni* subsp. jejuni 81-176 GenBank : AAD5138 4.1 | 5'-SalI-*pglA*$^{Cj}$ -SphI-3' | pNW1 923 | GTCGACAAGGAGGTGATCATT<br>AAAAATGAGAATAGGATTTTTA<br>TCACATGCAGGAGCAAGTATT<br>TATCATTTTAGAATGCCTATTA<br>TAAAAGCATTAAAAGATAGAAA<br>AGATGAAGTTTTTGTTATAGTG<br>CCACAAGATGAATACACGCAA<br>AAACTTAGAGATCTTGGCTTA<br>AAAGTAATTGTTTATGAGCTTT<br>CAAGAGCTAGTTTAAATCCTTT<br>TGTGGTTTTAAAGAATTTTTTT<br>TATCTTGCTAAGGTTTTGAAAA<br>ATTTAAATCTTGATCTTATTCA<br>AAGTGCGGCACACAAAAGCAA<br>TACCTTTGGAATTTTAGCAGC<br>AAAATGGGCAAAAATTCCTTAT<br>CGTTTTGCCTTAGTAGAAGGC<br>TTGGGATCTTTTTATATAGATC<br>AAGGTTTTAAGGCAAATTTAGT<br>GCGTTTTGTTATTAATAATCTT<br>TATAAATTAGGTTTTAAATTTG<br>CACACCAATTTATTTTTGTCAA<br>TGAAAGTAATGCTGAGTTTAT<br>GCGGAATTTAGGATTTAAGGA<br>AAGTAAAATTTGCGTGATAAAA<br>TCTGTAGGGATCAATTTAAAAA<br>AATTTTTTCCTATTTATGTAGA<br>ATCGGAAAAAAAAGAGCTTTT<br>TTGGAAAAAATTAAACATAGAT<br>AAAAAGCCCATTGTGCTTATG<br>ATAGCAAGAGCTTTATGGCAT<br>AAAGGTGTAAAAGAATTTTATG<br>AAAGTGCTACTATGCTAAAAG<br>ACAAAGCAAATTTTGTTTTAGT<br>TGGTGGAAGAGATGAAAATCC<br>TTCTTGTGCGAGTTTGGAGTT<br>TTTAAACTCGGGTGTGGTGCA<br>TTATTTGGGTGCTAGAAGTGA<br>TATAGTCGAGCTTTTGCAAAAT<br>TGTGATATTTTTGTTTTACCAA<br>GCTATAAAGAAGGCTTTCCTG<br>TAAGTGTTTTGGAGGCAAAAG<br>CTTGTGGCAAGGCTATAGTGG<br>TGAGTGATTGTGAAGGTTGTG<br>TAGAGGCTATTTCTAATGCTTA<br>TGATGGACTTTGGGCAAAAAC<br>AAAAAATGCTAAGGATTTAAG<br>CGAAAAAATTTCACTTTTATTA<br>GAAGATGAAAAATTAAGATTAA<br>ATTTAGCTAAAAATGCTGCCC<br>AAGATGCTTTACAATACGATG<br>AAAATAATATCGCACAGCGTT<br>ATTTAAAACTTTATGATAGGGT<br>AATTAAGAATGTATTGAGCATG<br>C | GTCGACAAGGAGGTGATCATTAAAAATGAGAATTG<br>GCTTTCTTTCACATGCTGGAGCCAGCATCTACCATT<br>TTAGAATGCCGATCATCAAAGCACTGAAAGATCGC<br>AAAGATGAAGTTTTTGTGATTGTCCCGCAAGATGAA<br>TATACACAGAAACTGAGAGATTTAGGCCTGAAAGTT<br>ATCGTGTATGAATTATCACGCGCAAGCCTGAATCC<br>GTTTGTTGTGCTTAAAAATTTCTTTTATCTTGCGAAA<br>GTGCTGAAAAACCTTAACCTGGATCTGATCCAATCT<br>GCAGCGCATAAATCAAACACATTTGGAATCCTGGC<br>TGCCAAATGGGCAAAAATCCCGTATAGATTTGCGC<br>TGGTCGAAGGCCTTGGATCTTTTTACATCGATCAA<br>GGCTTTAAAGCGAATCTGGTTCGCTTTGTGATCAA<br>CAACCTGTACAAACTTGGATTTAAATTTGCTCATCA<br>GTTTATCTTTGTT<br>AACGAATCAAACGCCGAATTTATGAGAAACCTGGG<br>CTTTAAAGAAAGCAAAATCTGCGTCATCAAATCAGT<br>TGGAATCAACCTGAAAAAATTTTTCCCGATCTATGT<br>GGAAAGCGAAAAGAAAGAACTGTTTTGGAAAAAAC<br>TGAACATCGATAAAAAACCGATTGTCCTGATGATCG<br>CACGCGCGCTTTGGCATAAAGGCGTTAAAGAATTT<br>TACGAAAGCGCTACAATGCTGAAAGATAAAGCCAA<br>CTTTGTCCTGGTTGGCGGAAGAGATGAAAACCCGT<br>CTTGTGCTTCATTAGAATTTCTGAATTCTGGCGTCG<br>TTCATTATCTTGGAGCCCGCTCAGATATTGTTGAAC<br>TGCTTCAAAATTGCGATATCTTTGTGCTGCCGTCAT<br>ATAAAGAAGGCTTTCCTGTGAGCGTCCTTGAAGCT<br>AAAGCCTGTGGCAAAGCGATTGTGGTCAGCGATTG<br>CGAAGGATGTGTTGAAGCAATCTCTAACGCGTATG<br>ATGGACTTTGGGCTAAAACAAAAAACGCCAAAGAT<br>CTGAGCGAAAAAATCTCTCTGCTGCTTGAAGATGA<br>AAAACTTCGCTTAAATCTGGCAAAAAACGCAGCGC<br>AAGATGCGCTGCAGTACGATGAAAACAACATCGCT<br>CAGAGATACCTTAAACTGTACGATCGCGTGATCAA<br>AAATGTCTAAGCATGC |
| *pglA*$^{Ng}$ | *Neisseria gonorrho eae* FA1090 GenBank : AAM157 78.1 | 5'-SalI-*pglA*$^{Ng}$ -SphI-3' | pNW1 932 | GTCGACAAGGAGGTGATCATT<br>AAAAATGAAAATCGTTTTTATC<br>ACAACAGTCGCATCCAGCATT<br>TACGGTTTCCGCGCCCCCGTC<br>ATTAAAAAATTAATCGGCAAAA<br>ACCATCAGGTGTATGCCTTTG<br>TATCGGAGTTTTCCGATAATG<br>AGTTGGACATTATCAGGGAAA<br>TGGGGGTTACACCCGTTACCT<br>ACCGGTCAAACCGCAGCGGG<br>GTAAACCCGTTTTCCGATATA<br>AAATCCACCTTCCTCATATTTA | GTCGACAAGGAGGTGATCATTAAAAATGAAAATTGT<br>GTTTATCACAACAGTCGCATCAAGCATTTATGGCTT<br>TCGCGCGCCGGTCATCAAAAAACTTATCGGAAAAA<br>ACCATCAAGTCTACGCATTTGTTTCTGAATTTTCAG<br>ATAACGAACTGGATATCATCAGAGAAATGGGCGTT<br>ACACCGGTGACATATAGAAGCAATCGCTCTGGAGT<br>TAACCCGTTTTCAGATATCAAAAGCACATTTCTGAT<br>CTTTAAAGCTCTGAAGAAAATTTCTCCGGATTTAGT<br>TTTTCCGTATTTTGCTAAACCGGTGATCTTTGGCAC<br>ATTTGCAGCGAAACTGGCCGGCGTGCCGCGCATT<br>GTCGGAATGCTTGAAGGCTTAGGATTTGCCTTTAC<br>ACCGCAGCCGGAAGGCATTCCGCTGAAAACAAAAA |

| | | | | | |
|---|---|---|---|---|---|
| | | | | AAGCACTCAAAAAAATATCGC<br>CGGATTTGGTTTTCCCTTATTT<br>CGCAAAACCCGTGATTTTCGG<br>CACTTTTGCCGCAAAATTGGC<br>AGGCGTGCCCAGAATCGTCG<br>GGATGCTGGAAGGTTTGGGAT<br>TCGCATTTACCCCGCAGCCGG<br>AAGGCATACCGTTAAAAACAA<br>AAATAATAAAGGGCATTTTGAT<br>TGCCCTGTACCGCATTGCCCT<br>GCCGATGTTGGAAAGCCTGAT<br>CGTATTAAACCCCGACGACAA<br>AGACGAGCTGCTGCATCAATA<br>CGGCATCAAAATAAAAAACAT<br>TCATATTTTGGGCGGAATCGG<br>TCTGGATTTGCGGCAATATCC<br>TTATTCCGAGGCGGATATTCC<br>CGATGAAAAAGAACCCGTAAA<br>ATTTCTCTTTATCGGCAGATTT<br>CTGAAAGAAAAGGGGATTGAT<br>GATTTTATTCGGGCGGCGGAA<br>CAGGTTAAGGGCAAATACCCC<br>GATACGGTTTTTACCGCTTTG<br>GGCGCAATCGACAAATCACGC<br>GGGGGGGGGGGAGATTTAGA<br>ACGCTTTATCGCCCGCGATAT<br>TATCCGTTTCCCCGGTTTTGT<br>GAACAATGTTTCCGAAGTGAT<br>AAAGGCGCATCATATATTCGT<br>ATTGCCGTCTTATTATAGGGA<br>AGGCGTTCCCCGAAGCACCC<br>AGGAGGCAATGGCCGTCGGC<br>AGGGCGGTGATTACGACGGA<br>TGTCCCCGGATGCAGGGAAA<br>CGGTTGCCGACAAGGTCAAC<br>GGCTTCCTGATCGAACCTTGG<br>AATCCCCGCATCTTGGCCGAA<br>AAAATGATTTATTTTATCGAAA<br>ACAGGGCTGCCGTCCGCCTG<br>ATGGCGAATGCAAGTTATGCG<br>ATTGCCAAAGATAAATTCGAT<br>GCCGAAAAAGTCGATTTGAAA<br>TTTCTCGATATTTTGAAGGCGT<br>AAGCATGC | TCATCAAAGGAATCCTGATCGCTCTTTATAGAATTG<br>CCTTACCGATGCTGGAATCACTTATCGTGTTAAATC<br>CGGATGATAAAGATGAACTGCTTCATCAATACGGC<br>ATCAAAATCAAAAACATCCATATCCTTGGCGGAATT<br>GGATTAGATCTGCGCCAGTATCCGTATAGCGAAGC<br>GGATATTCCGGATGAAAAAGAA<br>CCGGTCAAATTTCTGTTTATCGGCCGCTTTCTGAAA<br>GAAAAAGGAATCGATGATTTTATCAGAGCTGCCGA<br>ACAAGTCAAAGGCAAATATCCGGATACAGTTTTTAC<br>AGCACTGGGAGCGATCGATAAAAGCAGAGGCGGA<br>GGCGGAGATCTTGAAAGATTTATTGCACGCGATAT<br>TATCAGATTTCCGGGCTTTGTCAACAACGTTTCTGA<br>AGTGATCAAAGCGCATCATATCTTTGTGTTACCGTC<br>TTATTATCGCGAAGGAGTCCCGAGATCAACACAGG<br>AAGCTATGGCCGTTGGCCGCGCTGTGATTACAACA<br>GATGTTCCGGGATGCAGAGAAACAGTCGCCGATAA<br>AGTTAACGGATTTCTTATCGAACCGTGGAACCCGA<br>GAATCCTGGCTGAAAAAATGATCTACTTTATCGAAA<br>ATCGCGCAGCGGTTAGACTGATGGCAAACGCGAG<br>CTATGCTATCGCCAAAGATAAATTTGATGCAGAAAA<br>AGTGGATCTGAAATTTCTTGATATTTTAAAAGCGTA<br>AGCATGC |
| *pglA*<br>*Nm* | *Neisseria meningitidis*<br>C311#3<br>GenBank:<br>QXZ29626.1 | 5'-<br>SalI-<br>*pglA*$^{Nm}$-<br>SphI-<br>3' | pNW1933 | GTCGACAAGGAGGTGATCATT<br>AAAAATGAAAATCGTTTTTATC<br>ACAACAGTCGCATCCAGCATT<br>TACGGTTTCCGCGCCCCCGTC<br>ATTAAAAAATTAATCGGCAAAA<br>ACCATCAGGTGTATGCCTTTG<br>TATCGGAGTTTTCCGACAATG<br>AATTGGATATTATCAGGGAAA<br>TGGGGGTTACACCCGTTACCT<br>ACCGTTCAAACCGCAGCGGG<br>CTGAACCCGTTTTCGGATATA<br>AAATCCACCTTCCTCATCTTTA<br>AAGAACTCAAAAAAATATCGC<br>CGGATTTGGTTTTCCCTTATTT<br>CGCAAAACCCGTGATTTTCGG<br>CACTTTTGCCGCAAAACTGGC<br>AGGCGTGCCCAGAATCGTCG<br>GGATGCTGGAAGGTTTGGGAT<br>TCGCATTTACCCCGCAGCCGG<br>AAGGCATACCGTTAAAAACAA<br>AAATCATAAAGGGGATTTTGA<br>TTGCCTTATACCGCATTGCCC<br>TGCCGATGTTGGAAAGCCTGA<br>TTGTATTAAACCCCGACGACA<br>AAGACGAACTGACGGACAAAT | GTCGACAAGGAGGTGATCATTAAAAATGAAAATTGT<br>GTTTATCACAACAGTCGCATCAAGCATTTATGGCTT<br>TCGCGCGCCGGTGATCAAAAAACTGATCGGAAAAA<br>ACCATCAAGTCTACGCTTTTGTTTCTGAATTTTCAG<br>ATAACGAACTGGATATCATCAGAGAAATGGGCGTT<br>ACACCGGTGACATATAGAAGCAATCGCTCTGGACT<br>GAACCCGTTTTCAGATATCAAAAGCACATTTTTAAT<br>CTTTAAAGAACTGAAGAAAATTTCTCCGGATCTGGT<br>TTTTCCGTATTTTGCTAAACCGGTGATCTTTGGCAC<br>ATTTGCAGCGAAACTTGCCGGCGTGCCGCGCATTG<br>TCGGAATGCTTGAAGGCTTAGGATTTGCATTTACAC<br>CGCAGCCGGAAGGCATCCCGCTTAAAACAAAAATC<br>ATCAAAGGAATCCTGATCGCACTTTATAGAATTGCG<br>TTACCGATGCTGGAATCACTTATCGTCTTAAATCCG<br>GATGATAAAGATGAACTTACAGATAAATACGGCATC<br>AAAATCAAAAACATTCATATCTTAGGCGGAATTGGA<br>TTAGATCTGCGCCAATATCCGTATAGCGAAGCGGA<br>TATTCCGGATGAAAAAGAA<br>CCGGTTAAATTTCTTTTTATCGGCCGCTTTTTAAAA<br>GAAAAAGGAATCGATGATTTTATCAGAGCTGCCGA<br>ACAGGTCAAAGATAAATACCCGGATACAGTTTTTAC<br>AGCTCTTGGAGCCATTGATAAAAGCAGAGGCGGAG<br>GCGGAGATCTGGAAAGACTTGCAGCGCGCGATATT<br>ATCAGATTTCCGGGCTTTGTCAACAACGTTTCTGAA<br>GTGATCAAAGAACATCATATCTTTGTGTTACCGTCT |

| | | | | | |
|---|---|---|---|---|---|
| | | | | ACGGCATCAAAATAAAAAACA<br>TCCATATTTTGGGCGGAATCG<br>GTCTGGATTTGCGGCAATATC<br>CTTATTCCGAGGCGGATATTC<br>CCGATGAAAAAGAACCCGTAA<br>AATTCCTCTTTATCGGCAGATT<br>TCTGAAAGAAAAGGGGATTGA<br>TGATTTTATTCGGGCGGCGGA<br>ACAGGTTAAGGACAAATACCC<br>CGATACGGTTTTTACCGCTTT<br>GGGCGCAATCGACAAATCAC<br>GCGGGGGGGGGGGCGATTTG<br>GAACGGCTTGCCGCCCGCGA<br>TATTATCCGTTTCCCCGGTTTT<br>GTGAACAATGTTTCCGAAGTG<br>ATAAAAGAACATCATATATTCG<br>TATTGCCGTCTTATTATAGGG<br>AAGGCGTTCCCCGAAGCACTC<br>AGGAGGCAATGGCCGTCGGC<br>AGGGCAGTGATTACGACGGAT<br>GTCCCCGGATGCAGGGAAAC<br>GGTCGCCGACAAGGTCAACG<br>GCTTCCTGATCGAGCCTTGGA<br>ATCCCCGCATCTTGGCCGAAA<br>AAATGATTTATTTTATCGAAAA<br>CAGGGAAGCCGTCCGCCTGA<br>TGGGGAATGCAAGTTATGCGA<br>TTGCCAAAGATAAATTCGATG<br>CCGAAAAAGTCGATTTGAAAT<br>TGCTCGATATTTTGAAGGCGT<br>AAGCATGC | TATTATCGCGAAGGAGTCCCGAGATCAACACAAGA<br>AGCTATGGCCGTTGGCCGCGCTGTGATTACAACAG<br>ATGTTCCGGGATGCAGAGAAACAGTCGCCGATAAA<br>GTTAACGGCTTTTTAATCGAACCGTGGAACCCGAG<br>AATCCTGGCAGAAAAAATGATCTACTTTATCGAAAA<br>CCGCGAAGCTGTTAGATTAATGGGA<br>AACGCCAGCTATGCAATCGCGAAAGATAAATTTGA<br>TGCAGAAAAAGTGGATCTGAAACTGCTTGATATTCT<br>TAAAGCGTAAGCATGC |

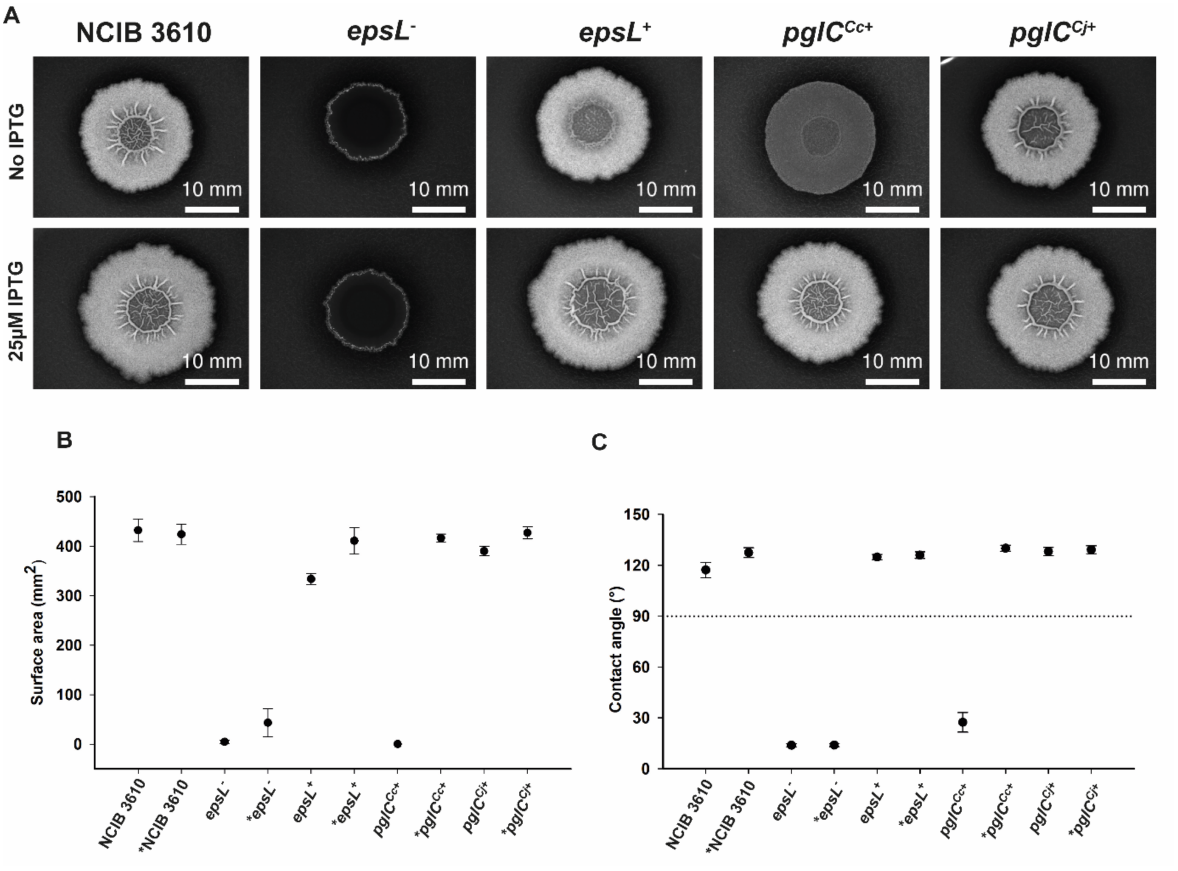

**Figure S7:** Colony biofilm morphology and hydrophobicity upon genetically complementing *ΔepsL-Bs* mutant with *pglC* of *Campylobacter*. (**A**) represents colony biofilm morphologies of wild-type (*B. subtilis* NCIB 3610), *ΔepsL* mutant (*epsL*$^{-}$ - NRS5907) and genetically complemented strains (*epsL*$^{+}$ - NRS5942, *pglC*$^{Cc+}$ - NRS6692, *pglC*$^{Cj+}$ - NRS6618). The colony biofilms were grown at 30 °C for 48 h under no IPTG and 25 µM IPTG-induced conditions. (**B**) represents the surface area calculated for the colony biofilm. (**C**) represents the respective sessile water drop analysis of the colony biofilms with a 5 µl water droplet on top. The representative images were taken after 5 minutes, except *epsL*$^{-}$ and *pglC*$^{Cc+}$ where the images were taken at 0 minutes due to the extreme hydrophilicity of the surface. Each data point in (**B**) and (**C**) represents the mean value for three biological replicates and their respective two technical replicates. Thus, error bars represent the standard deviation of six replicates. The dotted horizontal reference line in (**C**) represents the 90° contact angle which is a cut-off value for the hydrophobicity of *B. subtilis* biofilm. The data labeled with * on the x-axis represents the values of the biofilm grown under 25 µM IPTG condition.

**Table S7.** UniProt accession numbers and associated protein sequences of glycosyl transferases (GT) used for sequence identity.

| | UniProt Accession number | Organism | Sequence | Protein (gene) |
|---|---|---|---|---|
| *Bs* EpsD | P71053 | *Bacillus subtilis* | MTKKILFCATVDYHFKAFHLPYFKWFKQMGWEVHVAANGQTKLPYVDEKFSIPIRRSPFDPQNLAVYRQLKKVIDTYEYDIVHCHTPVGGVLARLAARQARRHGTKVLYTAHGFHFCKGAPMKNWLLYYPVEKWLSAYTDCLITINEEDYIRAKGLQRPGGRTQKIHGIGVNTERFRPVSPIEQQRLREKHGFREDDFILVYPAELNLNKNQKQLIEAAALLKEKIPSLRLVFAGEGAMEHTYQTLAEKLGASAHVCFYGFCSDIHELIQLADVSVASSIREGLGMNVLEGMAAEQPAIATDNRGHREIIRDGENGFLIKIGDSAAFARRIEQLYHKPELCRKLGQEGRKTALRFSEARTVEEMADIYSAYMDMDTKEKSV | Putative glycosyltransferase (*epsD*) |
| *Cc* PglA | A7ZET5 | *Campylobacter concisus* | MARIGFLSHADMSIHFFRRPIMQALKDMGHEVFAIAPKGNFTNELAKSFHTVTYELDKASLNPLTVINNSKKLSQILGELNLDLLQTGAHKSNVFGTFAAKNAGIKHVINLVEGLGSFYIDDDIKTKAVRFVMESLYKLSFAKADACIFVNDADPDYLISRNLIDKSKVYRIKSVGVDTAKFDPAITQAADLGEKKVILMIARAMWHKGVREFYEAAEILNGYKNCEFVFVGEGFAGNKSTADESFLKGGKVRYLGARNDIPQLLKASYLLALPSYKEGFPRTVLEAMSMAKAVVASDVTGCNEAVKDGYNGLLCKVKDASDLASKIKILLDDEALCAKLGANGRDWAVSEFDEKQIAKRYIEIYRKFIDV | *N, N'*-diacetylbacillosaminyl-diphospho-undecaprenol alpha-1,3-*N*-acetylgalactosaminyl transferase (*pglA*) |

| | | | | |
|---|---|---|---|---|
| *Cj* PglA | A0A2U0QT38 | *Campylobacter jejuni* | MRIGFLSHAGASIYHFRMPIIKALKDRKDEVFVIVPQDEYTQKLRDLGLKVIVYEFSRASLNPFVVLKNFFYLAKVLKNLNLDFIQSAAHKSNTFGILAAKWAKIPYRFALVEGLGSFYIDQGFKANLVRFVINSLYKLSFKFAHQFIFVNESNAEFMRNLGLKENKICVIKSVGINLKKFFPIYVESEKKELFWKNLNIDKKPIVLMIARALWHKGVKEFYESATMLKDKANFVLVGGRDENPSCASLEFLNSGAVHYLGARSDIVELLQNCDIFVLPSYKEGFPVSVLEAKACGKAIVVSDCEGCVEAISNAYDGLWAKTKNAKDLSEKISLLLEDEKLRLNLAKNAAQDALQYDENIIAQRYLKLYDRVIKNV | *N, N'*-diacetylbacillosaminyl-diphospho-undecaprenol alpha-1,3-*N*-acetylgalactosaminyl transferase |
| *Ng* PglA | Q5F602 | *Neisseria gonorrhoeae* | MKIVFITTVASSIYGFRAPVIKKLIGKNHQVYAFVSEFSDNELDIIREMGVTPVTYRSNRSGVNPFSDIKSTFLIFKALKKISPDLVFPYFAKPVIFGTFAAKLAGVPRIVGMLEGLGFAFTPQPEGIPLKTKIIKGILIALYRIALPMLESLIVLNPDDKDELLHQYGIKIKNIHILGGIGLDLRQYPYSEADIPDEKEPVKFLFIGRFLKEKGIDDFIRAAEQVKGKYPDTVFTALGAIDKSRGGGGDLERFIARDIIRFPGFVNNVSEVIKAHHIFVLPSYYREGVPRSTQEAMAVGRAVITTDVPGCRETVADKVNGFLIEPWNPRILAEKMIYFIENRAAVRLMANASYAIAKDKFDAEKVDLKFLDILKA | Glycosyl transferase family 1 (NGO_1765) |
| *Nm* PglA | Q9K1D9 | *Neisseria meningitidis* | MKIVFITTVASSIYGFRAPVIKKLIGKNHQVYAFVSEFSDNELDIIREMGVTPVTYRSNRSGLNPFSDIKSTFLIFKELKKISPDLVFPYFAKPVIFGTFAAKLAGVPRIVGMLEGLGFAFTPQPEGIPLKTKIIKGILIALYRIALPMLESLIVLNPDDKDELTDKYGIKIKNIHILGGIGLDLRQYPYSEADIPDEKEPVKFLFIGRFLKEKGIDDFIRAAEQVKDKYPDTVFTALGAIDKSRGGGGDLERLAARDIIRFPGFVNNVSEVIKEHHIFVLPSYYREGVPRSTQEAMAVGRAVITTDVPGCRETVADKVNGFLIEPWNPRILAEKMIYFIENREAVRLMGNASYAIAKDKFDAEKVDLKLLDILKA | Glycosyltransferase (*pglA*) |
| *Bs* EpsF | P71055 | *Bacillus subtilis* | MNSSQKRVLHVLSGMNRGGAETMVMNLYRKMDKSKVQFDFLTYRNDPCAYDEEILSLGGRLFYVPSIGQSNPLTFVRNVRNAIKENGPFSAVHAHTDFQTGFIALAARLAGVPVRVCHSHNTSWKTGFNWKDRLQLLVFRRLILANATALCACGEDAGRFLFGQSNMERERVHLLPNGIDLELFAPNGQAADEEKAARGIAADRLIIGHVARFHEVKNHAFLLKLAAHLKERGIRFQLVLAGDGPLCGEIEEEARQQNLLSDVLFLGTEERIHELMRTFDVFVMPSLYEGLPVVLVEAQASGLPCIISDSITEKVDAGLGLVTRLSLSEPISVWAETIARAAAAGRPKREFIKETLAQLGYDAQQNVGALLNVYNISTEKDHNR | Putative glycosyltransferase (*epsF)* |

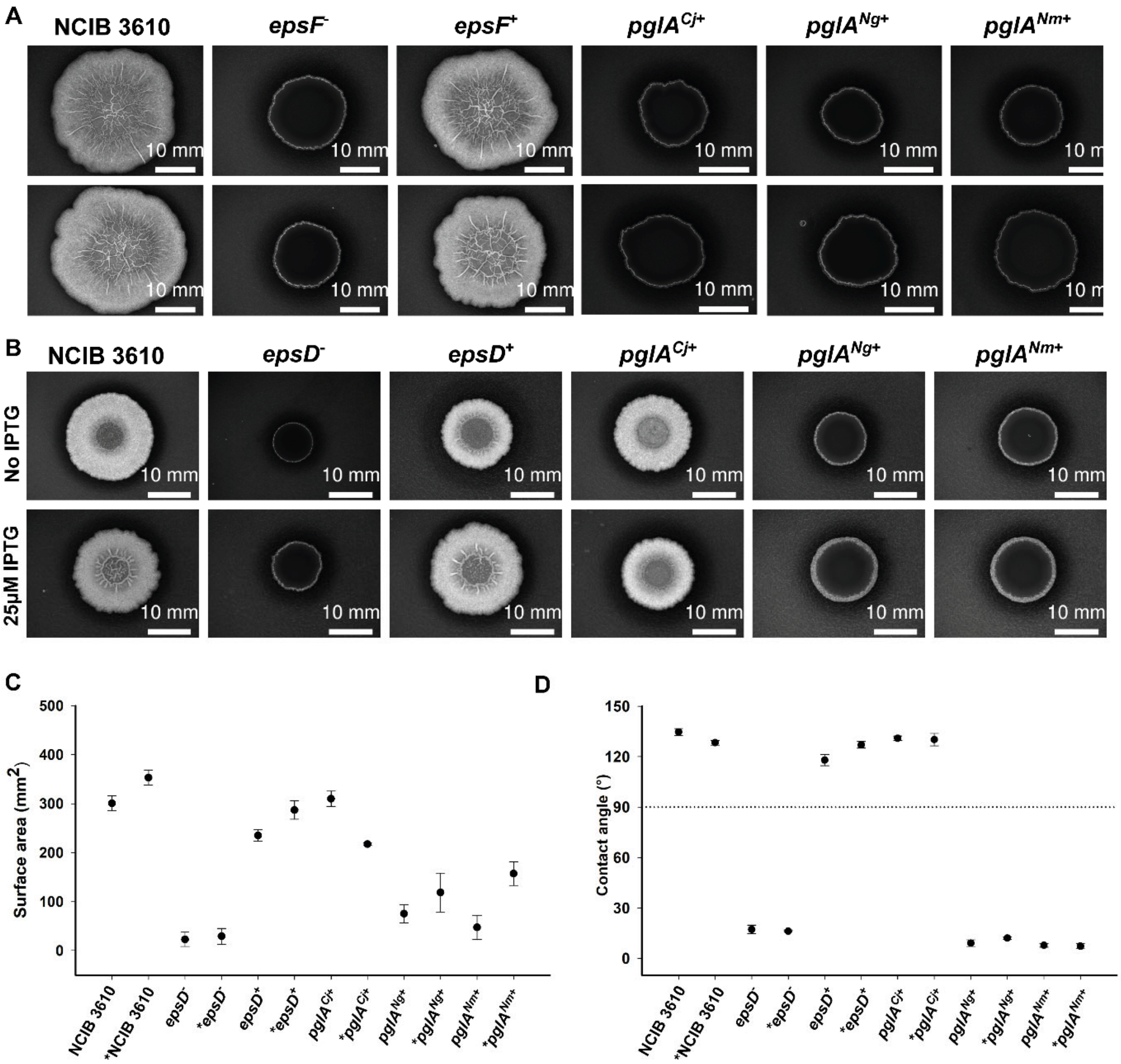


**Figure S8:** Colony biofilm morphology and hydrophobicity upon genetically complementing *ΔepsF-Bs ΔepsD-Bs* mutant with *pglA* of Campylobacter and Neisseria. (**A**) represents colony biofilm morphologies wild-type (*B. subtilis* NCIB 3610), *ΔepsF* mutant (*epsF*⁻ - NRS5904) and genetically complemented strains (*epsF*⁺ - NRS5961, *pglA*$^{Cj+}$ - NRS6628, *pglA*$^{Ng+}$ - NRS6629, *pglA*$^{Nm+}$ - NRS6630) (**B**) represents colony biofilm morphologies wild-type (*B. subtilis* NCIB 3610), *ΔepsD* mutant (*epsD*⁻ - NRS5905) and genetically complemented strains (*epsD*⁺ - NRS5930, *pglA*$^{Cj+}$ - NRS6605, *pglA*$^{Ng+}$ - NRS6619, *pglA*$^{Nm+}$ - NRS6620). The colony biofilms were grown at 30 °C for 48 h under no IPTG and 25 µM IPTG-induced conditions. (**C**) represents the surface area calculated for the colony biofilm. (**D**) represents the respective sessile water drop analysis of the colony biofilms with a 5 µl water droplet on top. The representative images of wild-type, *epsD*⁺ and *pglA*$^{Cj+}$ were taken after 5 minutes, whereas the images of *epsD*⁻ mutant, *pglA*$^{Ng+}$ and *pglA*$^{Nm+}$ were taken at 0 minutes due to extreme hydrophilicity of the surface in absence of biofilm (**C**) and (**D**) represent the mean value for three biological replicates and their respective two technical replicates. Thus, error bars represent the standard deviation of six replicates. The dotted horizontal reference line in (**D**) represents the 90°

contact angle which is a cut-off value for the hydrophobicity of *B. subtilis* biofilm. The data labeled with * on the x-axis represent the values of the biofilm grown under 25 μM IPTG condition.

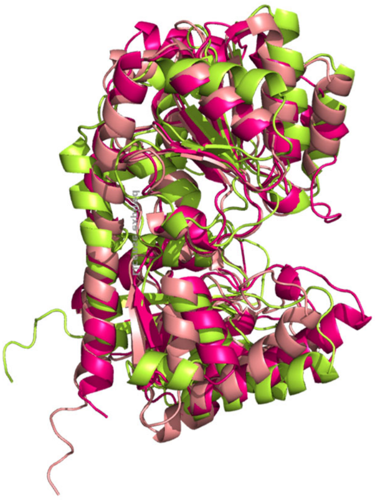

**Figure S9:** Structural alignment of PglA of *C. jejuni* (fluorescent green), *Bs* EpsD (Salmon pink) and *Bs* EpsF (dark pink). The superimposed structural model suggests that EpsF is structurally slightly different and EpsD aligns better with PglA than EpsF.

Protein sequence of EpsD-$His_6$ from *Bacillus subtilis (Bs):*

MTKKILFCATVDYHFKAFHLPYFKWFKQMGWEVHVAANGQTKLPYVDEKFSIPIRRSP
FDPQNLAVYRQLKKVIDTYEYDIVHCHTPVGGVLARLAARQARRHGTKVLYTAHGFHF
CKGAPMKNWLLYYPVEKWLSAYTDCLITINEEDYIRAKGLQRPGGRTQKIHGIGVNTER
FRPVSPIEQQRLREKHGFREDDFILVYPAELNLNKNQKQLIEAAALLKEKIPSLRLVFAG
EGAMEHTYQTLAEKLGASAHVCFYGFCSDIHELIQLADVSVASSIREGLGMNVLEGMA
AEQPAIATDNRGHREIIRDGENGFLIKIGDSAAFARRIEQLYHKPELCRKLGQEGRKTAL
RFSEARTVEEMADIYSAYMDMDTKEKSVHHHHHH

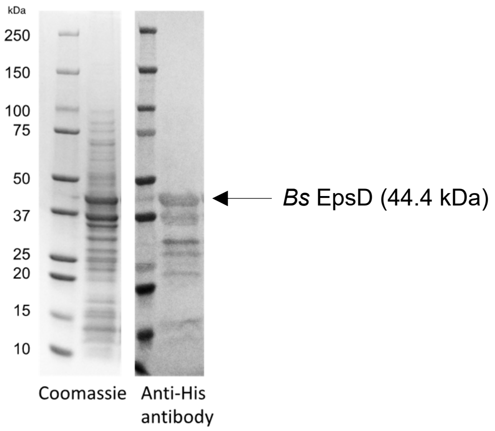


**Figure S10.** SDS PAGE (left panel) and Western blotting analysis (right panel) of *Bs* EpsD cell envelope fraction.

**Purification of EpsD.** The CEF of EpsD in 5.5 mL of 50 mM HEPES pH 7.5, 100 mM NaCl, was solubilized into 0.09% Triton X-100 (Anapoe-X-100) and incubated with rotation at 4 ºC for 2 h. The detergent-homogenized sample was centrifuged at 42,000 rpm for 65 minutes using a 70-Ti rotor. The supernatant was incubated with 1 mL Ni-NTA resin for 1 hour at 4 ºC. The resin was washed with 10 mL wash I buffer (50 mM HEPES pH 7.5, 100 mM NaCl, 15 mM imidazole, 5% glycerol, 0.09% Triton X-100, followed by a wash with 10 mL wash II buffer (50 mM HEPES pH 7.5, 100 mM NaCl, 45 mM imidazole, 5% glycerol, 0.09% TritonX-100). EpsD was eluted in 2 x 0.5 mL fractions of elution buffer (50 mM HEPES pH 7.5, 100 mM NaCl, 500 mM imidazole, 5% glycerol, 0.09% TritonX-100) and immediately desalted using a 5 mL desalting column in 50 mM HEPES pH 7.5, 100 mM NaCl, 5% glycerol, 0.09% TritonX-100. Desalted protein was eluted in 0.5 mL fractions from 1.5 mL – 3.0 mL. Fractions were pooled, flash-frozen in $LN_2$, and stored at –80 °C.

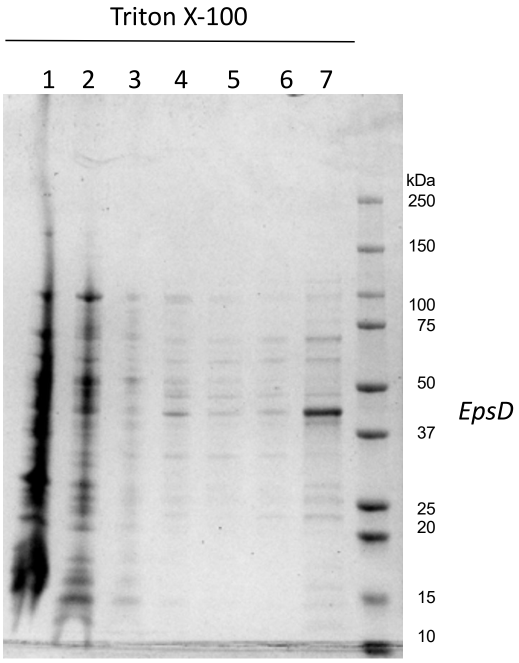


**Figure S11.** *Bs* EpsD detergent solubilization with Triton X-100 visualized by SDS PAGE (Coomassie). Lanes: 1) CEF; 2) flow through after solubilization and binding to Ni-NTA resin; 3) 45 mM imidazole wash; 4) 75 mM imidazole wash; 5) 100 mM imidazole wash; 6) 200 mM imidazole wash; and 7) desalted pure fraction from 400 mM imidazole elution.

**Und-PP-diNAcBac enzymatic synthesis (*Campylobacter concisus* PglC).** The Und-PP-Bac reaction was set up in a 7 mL scintillation vial. The reaction contained a total volume of 400 µL and consisted of 25 µM UndP, 50 µM UDP-diNAcBac, 100 nM *Cc* PglC, 50 mM HEPES pH 7.5, 100 mM NaCl, 0.1% Triton X-100 and 5 mM $MgCl_2$. The reaction contained a final concentration of 10% DMSO. The reaction was initiated by the addition of UDP-diNAcBac and allowed to proceed at ambient temperature for 30 minutes. The reaction was quenched with 2 mL of (2:1) $CHCl_3$/MeOH. The organic layer was washed three times with 500 µL PSUP (Pure Solvent Upper Phase = 15 mL $CHCl_3$, 240 mL MeOH, 1.83 g KCl, 235 mL $H_2O$) and concentrated under a stream of $N_2$. The crude oil was passed through a mini $Na_2SO_4$ pipette column to remove any remaining water and the eluted mixture was concentrated under $N_2$. The oil was then re-suspended in a mixture of (7:1) $CHCl_3$/MeOH and loaded on a silica column (column volume (CV) ~0.25 mL). The crude product was separated using a mobile phase gradient of 4 CV (7:1) $CHCl_3$/MeOH, 4 CV (5:1) $CHCl_3$/MeOH, and lastly 6 CV of 100% MeOH. Each fraction (~0.25 mL) was analyzed by TLC (solvent: 65:25:4 $CHCl_3$/MeOH/$H_2O$) and visualized with CAM staining

(0.5 g ceric ammonium sulfate, 12 g ammonium molybdate, 15 mL $H_2SO_4$, 235 mL $H_2O$). Subsequently, each fraction was quantified by the UDP-Glo biochemical assay *(vide infra).*

**UD-Glo biochemical assays to quantify Und-PP-diNAcBac.** Und-PP-diNAcBac concentration determination assays were performed with *Cc* PglA using the Promega UDP-Glo kit from Promega, which detects UDP generated over the course of the reaction. The quenching solution was prepared as described by Promega. A UDP-Glo standard curve was obtained using final [UDP] concentrations of 10 µM, 5 µM, 2.5 µM, 1.25 µM, 0.625 µM, 0.3125 µM, and 0.15625 µM from 10x UDP stocks in $H_2O$. The standard curve contained 10% DMSO. The PglA assays contained 100 nM *Cc* PglA, 0.1% Triton X-100, 50 mM HEPES at pH 7.5, 100 mM NaCl, 5 mM $MgCl_2$, 25 µM UDP-GalNAc, and Und-PP-diNAcBac in a final volume of 11 µL. An aliquot (5 µL) of each fraction from the Und-PP-diNAcBac purification (*vide supra*) was placed in a 1.7 mL Eppendorf and concentrated using the SpeedVac Vacuum Concentrator (10 minutes). Each concentrated Und-PP-diNAcBac fraction was resuspended in 1.1 µL of DMSO followed by the addition of assay buffer (7.7 µL). Then *Cc* PglA (1.1 µL of 1 µM) was added to the reaction mixture lacking the UDP-sugar for 2 minutes at ambient temperature. The reactions were initiated by the addition of UDP-GalNAc (1.1 µL of 250 µM in $H_2O$) and quenched with 11 µL of the UDP detection reagent after 30 minutes. The reaction mixture (20 µL) from each sample was transferred to a 96-well plate (white, nonbinding surface, Corning). The plate was shaken at low speed for 30 s and incubated for 1 h at 25 ºC, and luminescence was read on the plate reader. All luminescence values were background subtracted before converting to UDP.

**Radioactivity-based biochemical assays on *Bs* EpsD cell envelope fraction (CEF)**
EpsD *(Bs)* substrate specificity was measured using a radioactively-labeled UDP-[$^3$H]-sugar panel and an extraction-based assay.(3) Enzymatic reactions contained 20 µM Und-PP-Bac (2.5 µL of 200 µM in DMSO), UDP-[$^3$H]-sugar (1 µL in $H_2O$),[1] and 4 mg/mL EpsD CEF (2 µL of 50 mg/mL *total protein*) in a final volume of 25 µL of assay buffer (19.5 µL of 50 mM HEPES pH 7.5, 100 mM NaCl, 0.1% Triton X-100, 5 mM $MgCl_2$). The assay

contained a final concentration of 10% DMSO. Und-PP-Bac and UDP-[$^3$H]-sugar were pre-incubated in assay buffer for 30 seconds and the reactions were initiated by the addition of EpsD (*Bs*) CEF. An aliquot of 15 µL was quenched into 1 mL of 2:1 $CHCl_3$/MeOH after 10 minutes. The organic layer was washed three times with 500 µL PSUP (Pure Solvent Upper Phase = 15 mL $CHCl_3$, 240 mL MeOH, 1.83 g KCl, 235 mL $H_2O$). The organic layer was mixed with 5 mL Opti-Fluor O scintillation cocktail (PerkinElmer) and the combined aqueous layers were mixed with 5 mL EcoLite Liquid Scintillation Cocktail (MP Biomedicals). All layers were analyzed on a Beckman Coulter LS6500 scintillation counting system with quench compensation. EpsD *(Bs)* activity is background subtracted and reported as both dpm in the organic layer, and percentage of dpm in the organic layer normalized to the total dpm per quenched point. Error bars represent biological triplicate and calculated with GraphPad Prism 8.

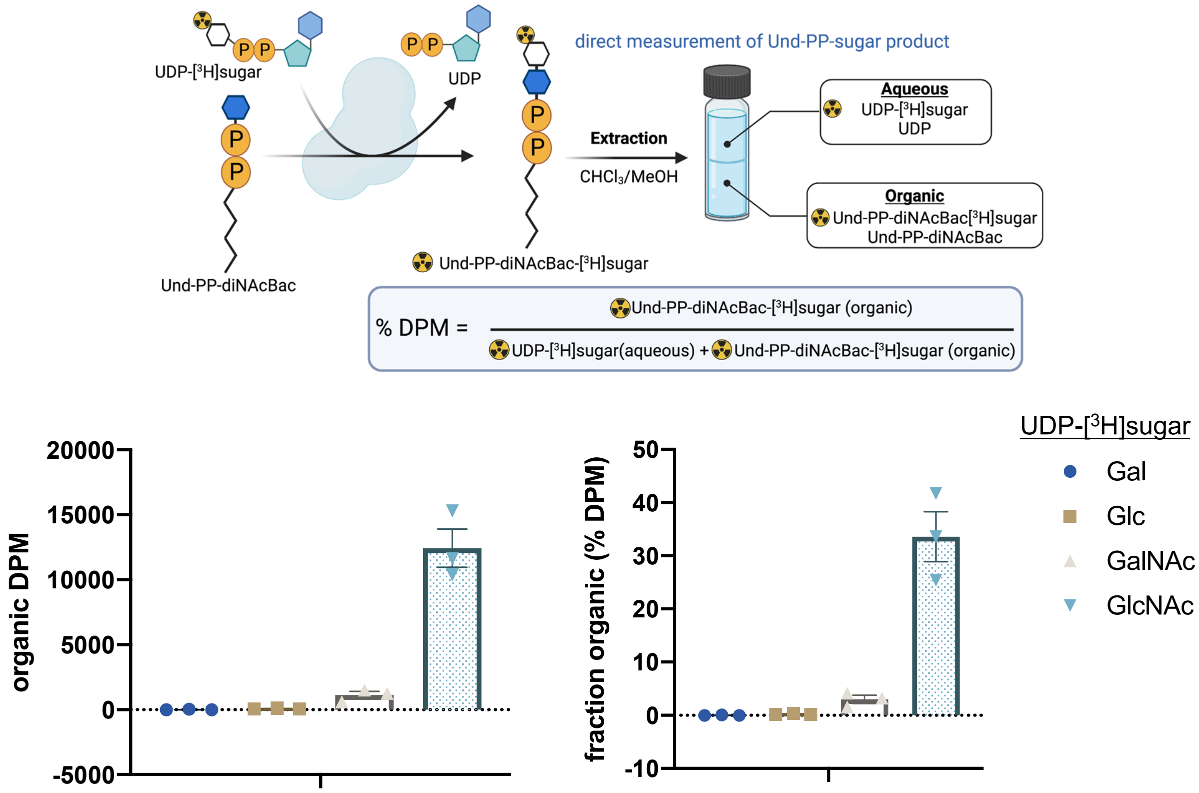


**Figure S12.** Substrate determination for the first glycosyl transferase in *B. subtilis,* EpsD, using a radioactivity-based assay. Error bars are given for mean ± SEM, n = 3.

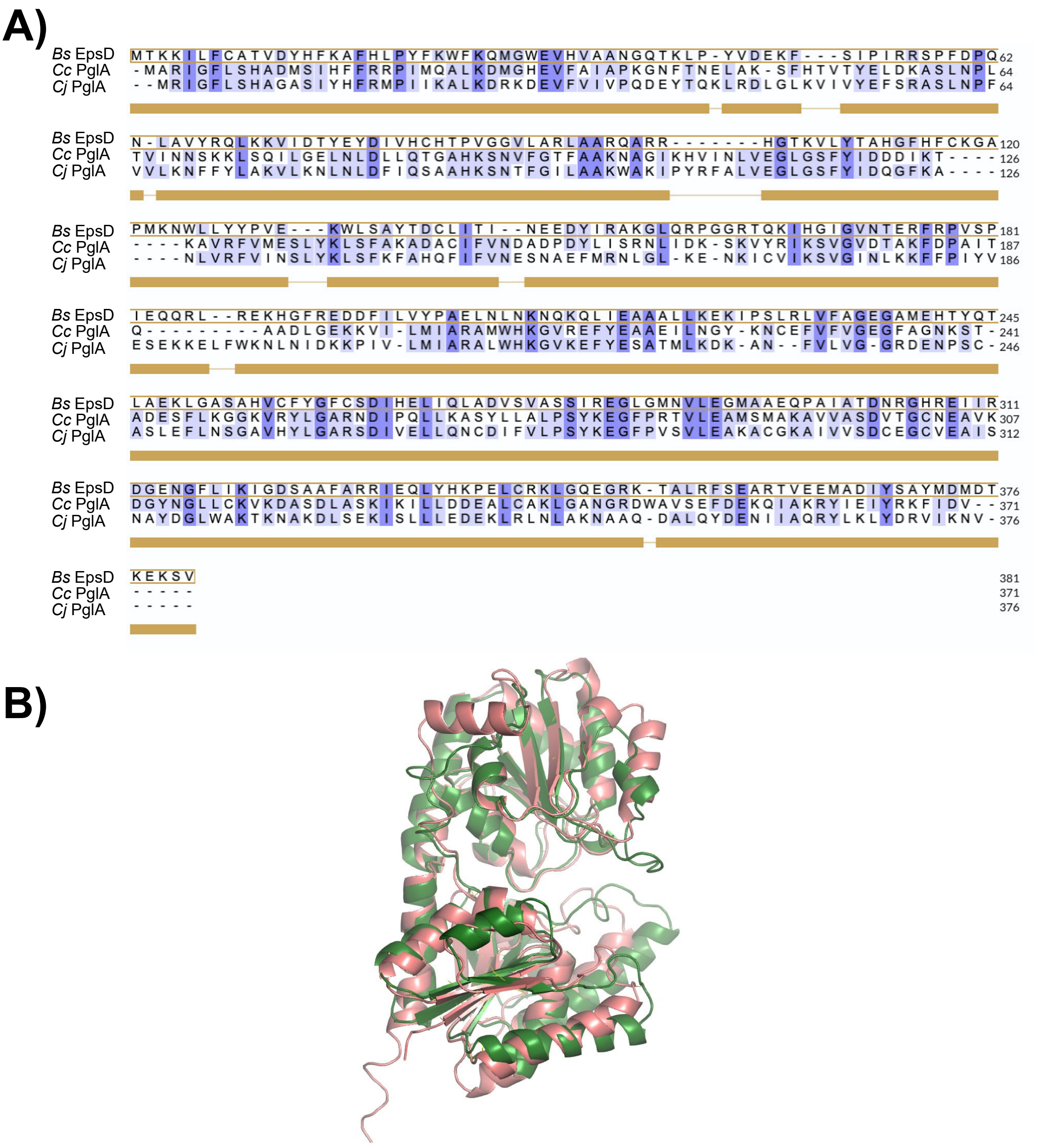


**Figure S13:** Comparison of *B. subtilis* EpsD (P71053) with PglAs from *C. concisus* (A7ZET5) and *C. jejuni* (A0A2U0QT38). **A)** Sequence alignment of EpsD of *B. subtilis* and PglA of *C. concisus* and *C. jejuni.* **B)** Superimposed AlphaFold models of *B. subtilis* EpsD (Salmon pink) and *C. concisus* PglA (green).
